# The human Origin Recognition Complex is essential for pre-RC assembly, mitosis and maintenance of nuclear structure

**DOI:** 10.1101/2020.08.06.240358

**Authors:** Hsiang-Chen Chou, Kuhulika Bhalla, Osama El Demerdesh, Olaf Klingbeil, Kaarina Hanington, Sergey Aganezov, Peter Andrews, Habeeb Alsudani, Kenneth Chang, Christopher R. Vakoc, Michael C. Schatz, W. Richard McCombie, Bruce Stillman

**Affiliations:** Cold Spring Harbor Laboratory, Cold Spring Harbor, New York 11746; Graduate Program in Molecular and Cellular Biology, Stony Brook University, Stony Brook, NY; Department of Computer Science, Whiting School of Engineering, John Hopkins University, Baltimore, Maryland 21218

**Keywords:** Initiation of DNA replication, Origin Recognition Complex, pre-Replicative Complex, CDC6, mitosis, CRISPR-Cas9 gene editing, nuclear structure.

## Abstract

The Origin Recognition Complex (ORC) cooperates with CDC6, MCM2-7, and CDT1 to form pre-RC complexes at origins of DNA replication. Here we report tiling-sgRNA CRISPR screens that show that each subunit of ORC and CDC6 are essential in human cells. Using an auxin-inducible degradation system, stable cell lines were created that ablate ORC2 rapidly, revealing multiple cell division cycle phenotypes. The primary defect in the absence of ORC2 was cells encountering difficulty in initiating DNA replication or progressing through the cell division cycle due to reduced MCM2-7 loading onto chromatin in G1 phase. The nuclei of ORC2 deficient cells were also large, with decompacted heterochromatin. Some ORC2 deficient cells that completed DNA replication entered into, but never exited mitosis. ORC1 knockout cells also demonstrated extremely slow cell proliferation and abnormal cell and nuclear morphology. Thus, ORC proteins and CDC6 are indispensable for normal cellular proliferation and contribute to nuclear organization.

## Introduction

Cell division requires the entire genome to be duplicated once and only once during the S-phase of the cell cycle, followed by segregation of the sister chromatids into two daughter cells. To ensure the complete and correct duplication of genomes, the initiation of DNA replication is highly regulated and begins with the assembly of a pre-Replication Complex (pre-RC) at origins of DNA replication throughout the genome (Bell and Labib, 2016). Among eukaryotes, studies in *Saccharomyces cerevisiae* have proved to be the best characterized system, from which individual proteins involved in DNA replication have been identified and studied extensively, including reconstitution with purified proteins of pre-RC assembly and the regulated initiation of DNA replication from pre-RCs (Evrin et al., 2009; Remus et al., 2009; Yeeles et al., 2015). In *S. cerevisiae*, pre-RC assembly begins with the hetero-hexameric Origin Recognition Complex (ORC), comprising Orc1-6 subunits, binding to each potential DNA replication origin (Bell et al., 1993; Bell and Labib, 2016; Bell and Stillman, 1992; Gibson et al., 2006). Chromatin-bound ORC then provides a platform for the assembly and recruitment for other pre-RC proteins. Cdc6 binds to ORC, followed by the binding of the Cdt1-Mcm2-7 complex to form head-to-head Mcm2-7 double hexamers to complete the formation of the pre-RC (Araki, 2011; Bell and Labib, 2016; Bleichert et al., 2017; Evrin et al., 2009; Heller et al., 2011; Remus et al., 2009). The Mcm2-7 double hexamer helicase precursor complex remains bound to DNA in an inactive state until it is activated by additional proteins and protein kinases (Bell and Labib, 2016). During S phase, Cyclin-dependent protein kinase (CDK) and the Cdc7-Dbf4-dependent protein kinase (DDK), Sld2, Mcm10, Dpb11, Sld3/7, DNA polymerase ε, Cdc45 and the GINS complex are recruited to activate MCM2-7 helicase (Araki, 2016; Araki et al., 1995; Bell and Labib, 2016; Kamimura, 2001; Kamimura et al., 1998; Takayama et al., 2003; Yeeles et al., 2015). The functional helicase consists of Cdc45-Mcm2-7-GINS (CMG) and when activated it unwinds the DNA in a bidirectional and temporally regulated manner from each origin (Bleichert et al., 2017).

In all eukaryote cells, including those from both yeast *S. cerevisiae* and human, each ORC1-5 subunit consists of a AAA+ or a AAA+-like domain and a winged helix domain (Bleichert et al., 2017; Chen et al., 2008; Li et al., 2018; Ocaña-Pallarès et al., 2020; Tocilj et al., 2017). In yeast, Orc1-6 remains as a stable complex bound to the chromatin throughout the cell division cycle (DePamphilis, 2003; Weinreich et al., 1999). ORC binds to A and B1 DNA sequence elements within the Autonomously Replicating Sequence (ARS), which contains a conserved ARS Consensus Sequence (ACS) (Bell and Labib, 2016; Bell and Stillman, 1992; Celniker et al., 1984; Deshpande and Newlon, 1992; Marahrens and Stillman, 1992; Rao and Stillman, 1995).

On the other hand, in human cells, there is no sequence-specific binding of ORC to the DNA, and the binding of ORC to chromosomes is dynamic (Vashee et al., 2001). ORC subunits do localize to specific sites within the chromosome, most likely via interactions with modified histones (Hossain and Stillman, 2016; Long et al., 2019; Miotto et al., 2016). One or more of the human ORC subunits dissociate from the complex soon after the pre-RC is formed. For example, ORC1 is ubiquitinated by the SCF^skp2^ ubiquitin ligase during the G1-S transition and then re-appears as cells enter mitosis (Kara et al., 2015; Kreitz et al., 2001; Méndez et al., 2002; Ohta et al., 2003). Human cell ORC1 is the first ORC subunit to bind to mitotic chromosomes and is inherited into the daughter cells where it recruits other ORC subunits and CDC6 to form the new pre-RCs (Kara et al., 2015; Okuno et al., 2001).

ORC is a conserved complex in eukaryotes, and it is essential for DNA replication in *S. cerevisiae, S. pombe, Xenopus and Drosophila*, since mutation or depletion of ORC prevents CDC6 binding and MCM loading onto DNA (Chuang et al., 2002; Pflumm and Botchan, 2001; Romanowski et al., 1996; Speck et al., 2005). Besides its function in the initiation of DNA replication, ORC protein subunits also play other important roles. In yeast, Orc1 directly interacts with silencing regulator Sir1 at the silent mating type loci to mediate transcriptional gene silencing and maintain heterochromatin (Bell et al., 1993; Fox et al., 1995; Hou et al., 2005; Triolo and Sternglanz, 1996). ORC1 also plays a role in transcriptional gene silencing in human cells (Hossain and Stillman, 2016). ORC2 depletion after pre-RC assembly resulted in spindle and DNA damage checkpoint activation, and impaired sister-chromatid cohesion (Shimada and Gasser, 2007). In *Drosophila*, Orc2 mutants showed reduced S phase cells, increased number of mitotic cells with abnormally condensed chromosomes and chromosome alignment defects, and more importantly, those mutants could not survive at late larval stage (Loupart et al., 2000; Pflumm and Botchan, 2001). In human, mutations in ORC1, ORC4, ORC6, CDT1, and CDC6 are detected in Meier-Gorlin syndrome patients (Bicknell et al., 2011b, 2011a; Guernsey et al., 2011; Hossain and Stillman, 2012; Munnik et al., 2015). ORC1 and ORC2 localize to centrosomes and ORC1 regulates the re-duplication of the centriole (Hemerly et al., 2009; Prasanth et al., 2004). ORC also localizes to telomeres via the TRF2 shelterin protein (Deng et al., 2009; Tatsumi et al., 2008). It was also shown that siRNA knockdown or CRISPR/Cas9 knockout of ORC1 resulted in loss of MCM2-7 from chromatin, abnormal duplication of centrioles, and a change in cell cycle stage distribution (Hemerly et al., 2009; Kara et al., 2015; McKinley and Cheeseman, 2017). ORC1, ORC2, ORC3, and ORC5 associate with heterochromatin, and depletion of any ORC subunits disrupt localization of heterochromatin and also causes abnormal heterochromatin decondensation in cells (Giri et al., 2016; Prasanth et al., 2010, 2004). ORC2 and ORC3 also specifically localize to centromeric heterochromatin during late S phase, G2 and mitosis and removal of these proteins causes decondensation of centromeric a-satellite (Craig et al., 2003; Prasanth et al., 2010, 2004).

There is an emerging debate, however, about the essential nature of ORC in human cells (Bell, 2017). ORC is overexpressed in numerous cancerous cell lines (McNairn and Gilbert, 2005) and HCT116 colorectal cancer cells can survive with only 10% of the ORC2 protein level (Dhar et al., 2001). More importantly, it was reported that HCT116 p53^-/-^ (*TP53^-/-^*, but we henceforth use *p53-/-*) cells in which expression of either ORC1 or ORC2 subunit was eliminated using CRISPR-Cas9 mediated gene ablation could still proliferate (Shibata et al., 2016). Here we developed a genetic method to address the function of the pre-RC proteins ORC and CDC6, particularly focusing on the ORC1 and ORC2 subunits. We demonstrate that ORC proteins are essential for normal cell proliferation and survival in human cells. Moreover, ORC1 or ORC2 depleted cells showed multiple defects in progression through cell division cycle, including DNA replication and mitosis, as well as defects in nuclear structure.

## Results

### ORC1-6 and CDC6 are essential for cell survival

Despite the implication that ORC in performs a critical role in DNA replication as well as cellular proliferation, there are conflicting observations about the essential role for the ORC1 and ORC2 subunits in human cells (Shibata et al., 2016). To further confound the issue, examination of the genes essential for cell survival in the DepMap database that summarize results from whole-genome CRISPR screens (GeCKO 19Q1 and Avana 20Q2 libraries) (Meyers et al., 2017; Tsherniak et al., 2017), show that ORC1 is found to be a common essential gene while ORC2 is listed as non-essential in tested cell types (Figure 1—Figure supplement 2a). Other members of the pre-RC proteins - ORC3-6, CDC6, MCM2-7 (data not shown) and CDT1 (data not shown) are all classified as common essential by at least one of the two whole genome CRISPR screens (GeCKO and Avana) but not necessarily both in DepMap.

**Figure 1.**
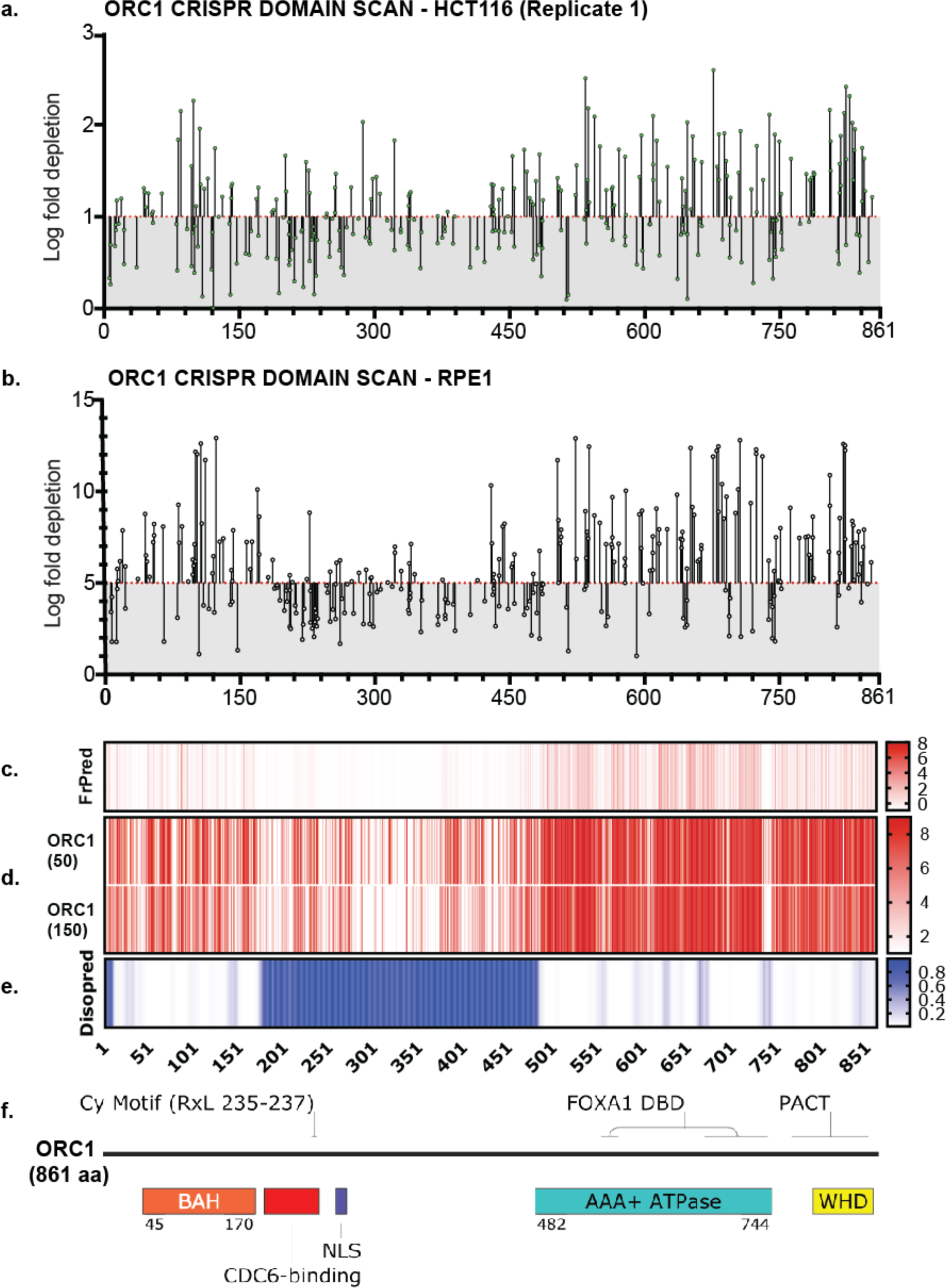
ORC1 is essential to HCT116 and RPE-1 cell lines. (a) tiling sgRNA map ORC1 (replicate 1) in HCT116. Mapped as Log fold depletion (inverted LFC scale) as calculated by MaGeCK (Li et al., 2014) on y axis vs the codon that is disrupted by the guide RNA on the x axis. Effect of guide RNA is interpreted as essential if its depletion is more than 1 log fold (red dotted line). (b) tiling sgRNA map of ORC1 in RPE-1. Effect of guide RNA is interpreted as essential if its depletion is more than 5 log fold (red dotted line). (c) FrPred (https://toolkit.tuebingen.mpg.de/frpred) of hORC1 (NP_004144.2) shown as gradient heat map of conservation score vs amino acid position. (d) Consurf (https://consurf.tau.ac.il/) of hORC1 – (upper) ORC1 (50) subset (50 HMMER Homologues collected from UNIREF90 database, Max sequence identity = 95%, Min sequence identity 50, Other parameters = default), and (lower) ORC1 (150) subset (150 HMMER Homologues collected from UNIREF90 database, Max sequence identity = 95%, Min sequence identity 35, Other parameters = default). Data represented as heat map of Conservation scores of each amino acid position. (e) Disopred (http://bioinf.cs.ucl.ac.uk/psipred/) plot of hORC1 – heat map representing amino acids within intrinsically disordered regions of the protein. (f) Schematic of domain architecture of ORC1.

To address the question of essentiality and to map functional domains within the ORC and CDC6 proteins, we used unbiased tiling-sgRNA CRISPR depletion screens which have been shown to be informative of not only the overall essentiality of the protein to cell fitness but also directly help nominate previously unknown, functionally essential regions in these in proteins, DNA and RNA (He et al., 2019; Hsu et al., 2018; Montalbano et al., 2017; Shi et al., 2015; Wang et al., 2019). We employed a tiling sgRNA CRISPR scheme using designed libraries of guide RNA targeting every possible PAM sequence 5′-NGG-3′ (S*treptococcus pyogenes* Cas9) across each exon of *ORC1-6* and *CDC6*. These CRISPR libraries also included control guide RNAs targeting either known core essential genes or those targeting non-essential gene loci or no loci at all (Miles et al., 2016). The total library comprised 882 guides targeting *ORC1-6* and *CDC6*, 1602 negative controls (Used in GeCKO V2 library - “NeGeCKO” (Sanjana et al., 2014), negative controls used in The Sabatini/Lander CRISPR pooled library (Park et al., 2016), Rosa26, CSHL in-house negatives (Lu et al., 2018; Tarumoto et al., 2018) and 43 positive controls (median of 3 guides targeting essential genes such as those encoding CDK1, CDK9, RPL9, PCNA etc.) (Supplement Table1_guides). Parallel screens were done in HCT116-Cas9 and RPE-1-Cas9 cell lines and the relative depletions of guide RNA in the cell populations between day 3 and day 21 were compared using the guide read counts generated by illumina based next-generation sequencing (n = 2 for HCT116, n = 1 for RPE-1) and the data was analyzed with MAGeCK (Li et al., 2014). The screens performed well as shown by the consistent log fold change (LFC) pattern of depletion or enrichment of positive and negative controls respectively - although the absolute values and range of LFCs were cell-line specific.

As expected, overall the CRISPR-Cas9 screen had a more profound effect on the diploid RPE-1 cells as compared to a transformed cell line like HCT116. The LFC threshold of ‘essentiality’ was set at the value at which a guide RNA was depleted more than every negative control as well as ≥ to the median depletion of guides targeting positive controls (Figure 1—Figure supplement 1b-d). In two replicate screens of HCT116, LFC ≥ -1 and LFC ≥ -5 in RPE-1, were found to be the cut-off for log fold depletion, above which the regions of the protein can be called essential. The fact that many such regions exist across the protein of course also imply the protein itself is essential to the cell line. Importantly, screens showed significant drop out of guide RNAs that target regions that have been previously annotated by structure and function (Figure 1a-b, 1f, Figure 2a-c, Figure 2—Figure supplement 1a, 3a-c, 3g, 4a-c, 4g, 4h-j, 4n, 5a-c, 5g, 5h-j, 5n). To visualize the tiling sgRNA data mapped to translated protein sequence in relation to its amino acid conservation and disorder we used NCBI RefSeq coding sequences (NP_004144.2, NP_006181.1, NP_862820.1, NP_859525.1, NP_002544.1, NP_055136.1, NP_001245.1) for three analyses - (1) FrPred (Adamczak et al., 2004; Fischer et al., 2008) (https://toolkit.tuebingen.mpg.de/frpred) server that calculates a conservation score based on amino acid variability as well as the probability of it being a functional ligand binding or catalytic site at each amino acid position of the input sequence (Figure 1c, Figure 2—Figure supplement 1b, 3d, 4d, 4k, 5d, 5k); (2) Consurf (Ashkenazy et al., 2016) (https://consurf.tau.ac.il/) server which analyses the probability of structural and functional conservation despite amino acid variability for any given position of input sequence (Figure 1d, Figure 2—Figure supplement 1c, 3e, 4e, 4l, 5e, 5l). We ran this server overall with default parameters except for the number of species to include. In one analysis we chose 50 representative species homologues with maximum and minimum percent identity set to 95 and 50 respectively, and in the other we increased the species to 150 and set max. and min. percent identity to 95 and 35 to compare a larger evolutionary subset. In both analyses the UNIREF90 database was used, which consists of cluster sequences that have at least 90% sequence identity with each other into a single UniRef entry, thus increasing the representative diversity of species considered in the output. And lastly, 3) Disopred tool (Buchan and Jones, 2019; Jones and Cozzetto, 2015) (http://bioinf.cs.ucl.ac.uk/psipred/) that scores for intrinsically disordered regions (IDRs) that are usually not well conserved yet found to be functionally essential in many proteins (Figure 1e, Figure 2—Figure supplement 1d, 3f, 4fm 4m, 5f, 5m). In this similar way, we also performed tiling sgRNA screens for MCM2-7 and CDT1 and found them to be essential, but including that data was currently beyond the scope of this work.

**Figure 2.**
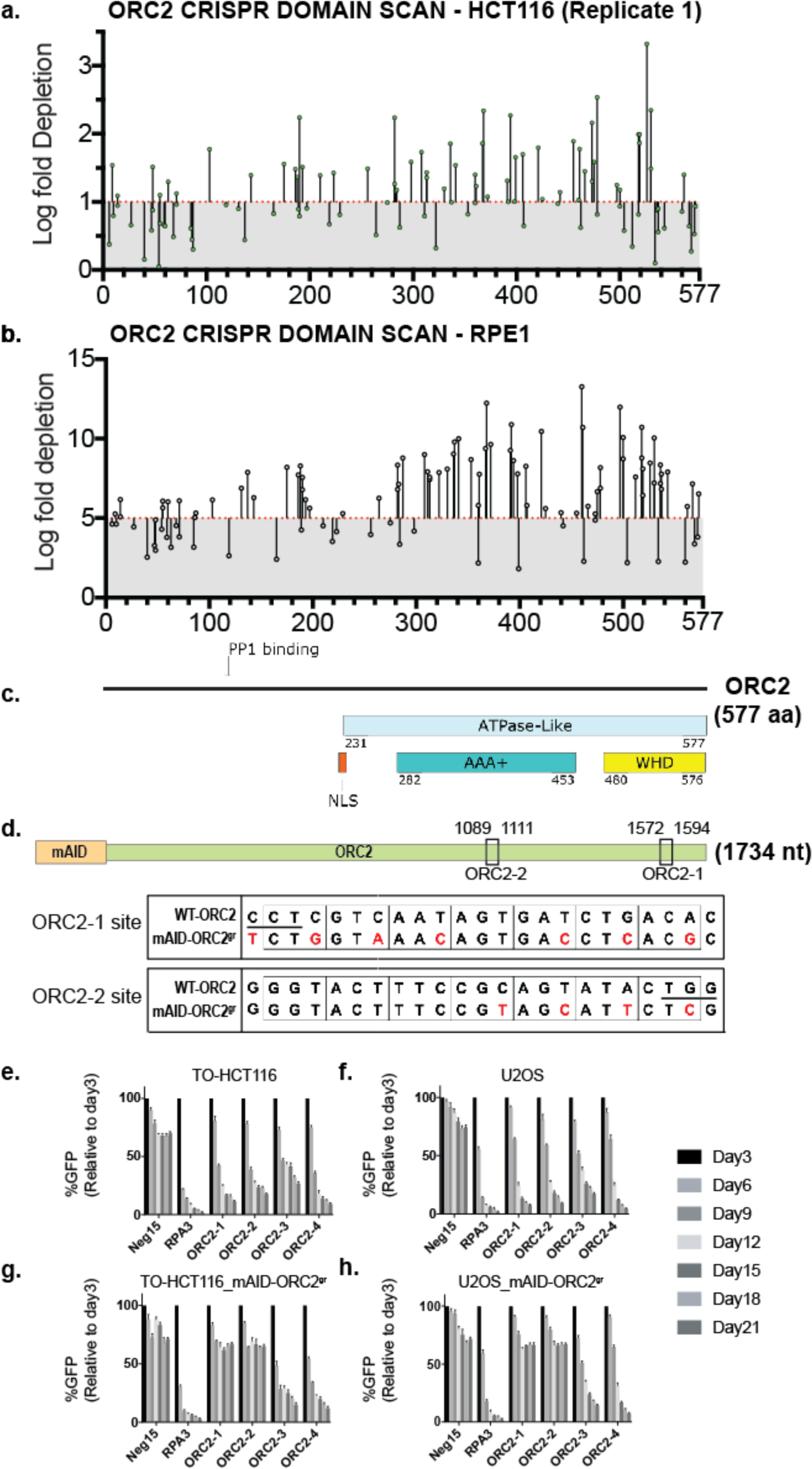
*ORC2* is essential in HCT116 and RPE-1 and both by tiling sgRNA and single guide CRISPR knock-down in presence of sgRNA-resistant mAID-ORC2^gr^. (a) tiling sgRNA map of ORC2 (replicate 1) in HCT116. Mapped as Log fold depletion (inverted LFC scale) as calculated by MaGeCK (Li et al., 2014) on y axis vs the codon that is disrupted by the guide RNA on the x axis. Effect of guide RNA is interpreted as essential if its depletion is more than 1 log fold (red dotted line). (b) tiling sgRNA map of *ORC2* for RPE-1. (c) schematic of ORC2 protein showing annotated structural or functional domains. (d) The top panel shows the mAID degron was fused to *ORC2* transgene at the N-terminus, and the two black boxes indicate ORC2-1 and ORC2-2 sgRNAs targeting regions. The numbers represent nucleotide positions in the *ORC2* cDNA. The lower two panels show the comparison of wild type and mAID-ORC2^gr^ with silent mutations at the indicated sgRNA targeting sites. The red color marks the mismatches. Protospacer-adjacent motif (PAM) site is underlined in the wild type sequence. (e-h) Negative-selection time course assay that plots the percentage of GFP positive cells over time following transduction with the indicated sgRNAs with Cas9. Experiments were performed in (e) TO-HCT116, (f) U2OS, (g) TO-HCT116_mAID-ORC2^gr^, and (h) U2OS_mAID-ORC2^gr^ cell lines. The GFP positive percentage was normalized to the Day3 measurement. N = 3. Error bars, mean ± SD.

For the purpose of this paper, we focus on discussing the tiling-sgRNA screens for *ORC1* and *ORC2* and compare the phenotypic differences with DepMap datasets as well as the previous study that classified these two subunits as non-essential. The pattern of depletion for *ORC1* showed that the strongest (higher in the plot) depletions map within the *B*romo-*A*djacent *H*omology (BAH) domain, *A*TPase *A*ssociated with diverse cellular *A*ctivities domain (AAA+) and the *W*inged-*H*elix *D*omain (WHD), all of which are crucial for its function (Figure 1a, 1b, 1f). The same is evident in both cell lines for *ORC2* where the highly depleted guides cluster to the ATPase-like C terminal end of ORC2 (Figure 2a-c). Interestingly, we found there were functionally essential regions in the lesser conserved IDR regions of both ORC1 and ORC2. Our lab has recently reported that a Cyclin-motif bearing region of ORC1 (180 - 240 aa) is essential for binding to CDC6 at the right time during the cell cycle to enable pre-RC formation (Hossain et al., 2019). And similarly, based on homology, a previously identified putative NLS motif (Lidonnici et al., 2004) also had hits nominating that region as essential in our *ORC1* screen (Figure 1f). There are other essential regions that correspond to novel protein binding sites that are separately being investigated.

When we compared DepMap CRISPR Achilles (Avana 20Q2 library) dataset and a combined RNAi dataset of cell lines, it indicated that using the CRISPR method, with a gene effect score of less than -0.5, *ORC1* classified as common essential in > 90 percent of the cell lines, while with RNAi datasets with that same cut-off, it classified as essential in only about 45 percent of the same cell lines (Figure 1—Figure supplement 2b-c). It is evident that the method of choice did have a bearing on the phenotypic outcome of the knock-down. The study by Shibata et al. (2016) that found *ORC1* and *ORC2* to be non-essential also used CRISPR editing as the method of knock-down, but also performed long term selection for cell proliferation to obtain *ORC1^-/-^* of *ORC2^-/-^* cells. We therefore determined our screen had guide RNAs that were used in either of the DepMap dataset or used in the directed study (Shibata et al., 2016). For *ORC1* and *ORC2* sgRNAs that were used in DepMap datasets, there was a variation in their phenotype as measured by LFC values, with some guides classifying *ORC1* and *ORC2* as essential and others not (Figure 2—Figure supplement 2a-c). Of note is the fact that the guide used to target *ORC2* in the Shibata et al. (2016) study showed activity very close to the cut-off in HCT116 cells and scored as non-essential in RPE-1. It is important to note that when a guide targeting a relatively non-essential region allows for the cells to proliferate, no conclusion can be made about the protein being essential. The Shibata et al. (2016) study used that single guide to insert a blasticidin and a poly A cassette into the locus, presumably disrupting its transcription significantly, while our single-guide-per-locus type of screen did not introduce such large insertions. We find that ORC1-6, CDC6 are essential in both cell lines tested, and that the depletion of these proteins negatively affected the diploid RPE-1 cells more. The results of these screens and the published DepMap conclusions, especially about *ORC2*, suggest that using too few guides to target proteins can lead to artifactual observations both in terms of essentiality or non-essentiality, and that overall gene-effect is affected by the combination of the choice of guide RNA and the cell line studied. At this point we selected guide RNAs that target *ORC2* from our tiling sgRNA screen to study them individually (Figure 2—Figure supplement 2d). We also received *ORC1* and *ORC2* deficient stable cell lines from the authors of the previous study (Shibata et al., 2016) for further analysis.

### Rapid ORC2 removal in cancer cells impedes cell growth and causes DNA damage

Depleting ORC2 with an siRNA approach was a slow process that took at least 24-48 hours, however, using this approach various phenotypes have been observed, including G1 arrest, an S-phase defect, and abnormally condensed chromosomes as well as defects in mitosis (Prasanth et al., 2010, 2004). These phenotypes can be an outcome of accumulated errors that happen during any phase of cell cycle and thus it is hard to distinguish between primary and secondary phenotypes associated with the loss of ORC2. Therefore, we used CRISPR/Cas9 in combination with tagging ORC2 with an auxin inducible degron (mAID) to construct cell lines in which the endogenous ORC2 was knocked out by CRISPR, and the complementing CRISPR-resistant mAID-ORC2 could be then rapidly removed from cells, allowing exploration of the importance of ORC2 at different stages of cell division cycle (Natsume et al., 2016; Nishimura et al., 2009). To mediate the endogenous ORC2 knockout, four sgRNAs were selected from the CRISPR screen to validate the screening result. For complementation, mAID-tagged sgRNA resistant ORC2 (mAID-ORC2^gr^) was constructed with the mAID degron fused to the ORC2 amino-terminus, and the *ORC2* cDNA was edited to harbor multiple mismatches based on two of the sgRNAs, ORC2-1 and ORC2-2 (Figure 2d). The mAID-ORC2^gr^ was transduced into the TO-HCT116 cell line, which expresses a doxycycline-inducible *Oryza sativa* (Asian rice) *TIR1* (OsTIR1) gene that encodes a plant auxin-binding receptor that interacts with the conserved E3 ubiquitin ligase SCF complex to degrade mAID-tagged proteins (Natsume et al., 2016; Nishimura et al., 2009), or the U2OS cell line to perform ORC2 depletion or genetic complementation, respectively. The effects of four sgRNAs were tested. In the negative-selection CRISPR/Cas9 experiment, cells expressing a positive control RPA3 sgRNA and all four ORC2 sgRNAs, but not the negative control Neg15 sgRNA, were outcompeted over 3 weeks in culture by non-transduced cells that lacked any sgRNA (Figure 2e-f). The CRISPR competition effects by ORC2-1 and ORC2-2 sgRNAs were rescued by mAID-ORC2^gr^ in both TO-HCT116_mAID-ORC2^gr^ and U2OS_mAID-ORC2^gr^ cell lines (Figure 2f-h).

To acquire cloned cells to study ORC2 depletion phenotypes, TO-HCT116_mAID-ORC2^gr^ cells were depleted of the *ORC2* gene with sgRNA ORC2_1 by the CRISPR/Cas9 editing technique then individual clones isolated after single cell sorting. Five cell lines, ORC2_H-1, ORC2_H-2, ORC2_H-3, ORC2_H-4, and ORC2_H-5, were obtained from two independent CRISPR/Cas9 knockout experiments done about 6 months apart. Sequencing of the target sites showed that the ORC2_H-1 and H-3 cell lines had heterozygous mutations at the sgRNA targeting site which create premature stop codons downstream of the target site (Figure 3—Figure supplement 1a). On the other hand, H-2, H-4, and H-5 are homozygous with an identical two-nucleotide-deletion, creating a nonsense mutation at the sgRNA targeting site. Although ORC2_1 sgRNA targets the C-terminus of ORC2, no truncated form of protein was detected by western blot. Our ORC2 rabbit polyclonal antibody was produced against amino-terminus half of the ORC2 protein. The LTR-driven mAID-ORC2^gr^ was low in expression compared to expression in the TO-HCT116, RPE-1 and IMR90 cells, but was sufficient to complement the loss of endogenous ORC2 done by CRISPR knockout (Figure 3a).

**Figure 3.**
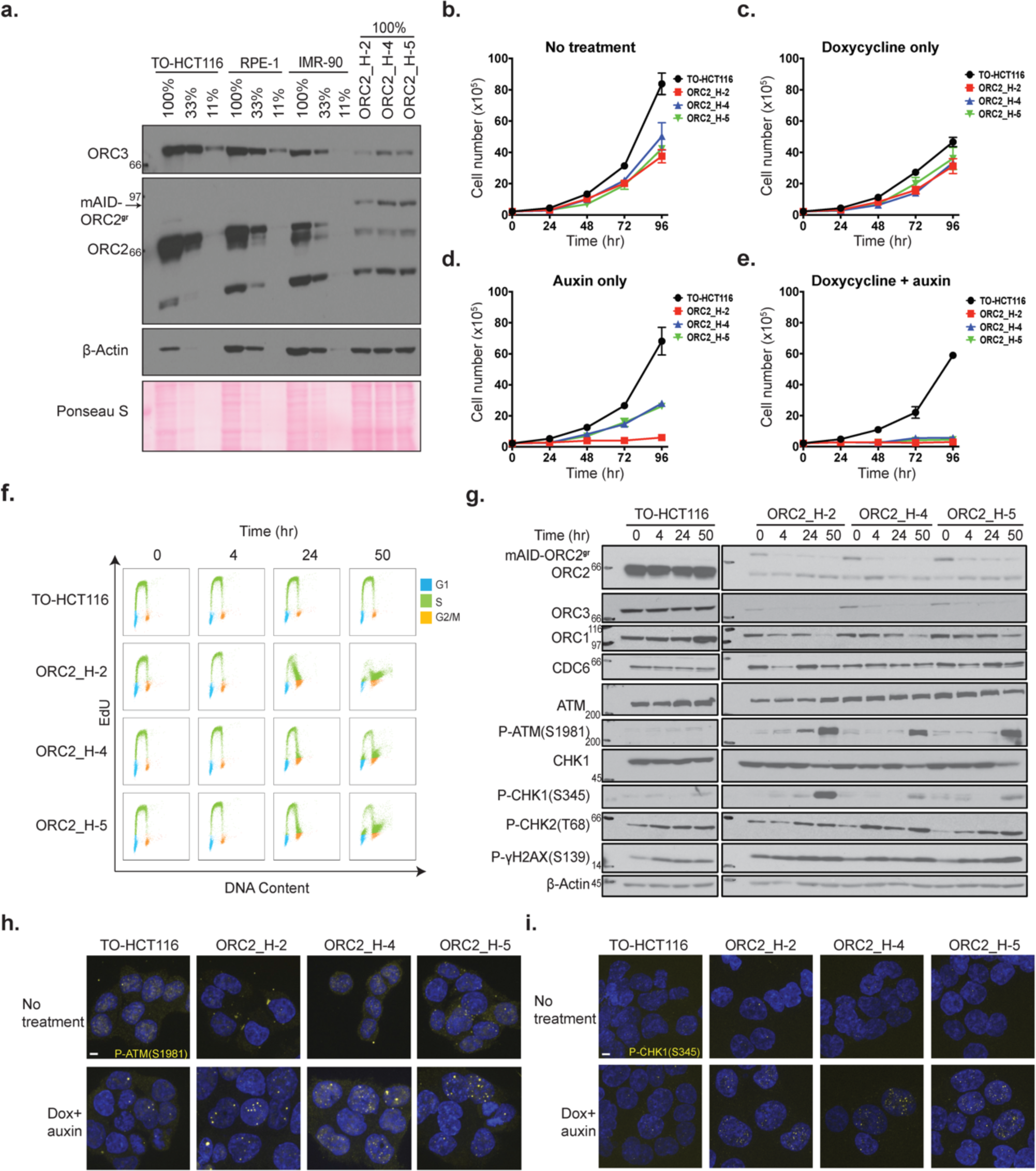
Characterization of CRISPR/Cas9 ORC2 knockout complemented with sgRNA resistant ORC2^gr^ cell lines. (a) Protein level of ORC2, mAID-ORC2^gr^, and ORC3 in human TO- HCT116, hTERT-RPE1, human diploid IMR-90 cell line, and three ORC2 KO cell lines. (b-e) Cell lines growth rates under (b) normal condition, (c) doxycycline only, (d) auxin only, and (e) Dox+auxin containing medium, respectively. The x axis indicates hours after adding doxycycline or auxin if any. The y axis reflects the cell number (x10^5^). N = 3 (biological repeats). Error bars, mean ± SD. (f) Cell cycle analysis of TO-HCT116, ORC2_H-2, ORC2_H-4, and ORC2_H-5 cell lines after mAID-ORC2^gr^ depletion. Cells were treated with 0.75 µg/ml of doxycycline for 24 hours before auxin treatment. Cells were pulse labeled with 10 μM EdU for 2 hours before harvesting at 0, 4, 24, and 50 hr. time points. The x axis indicates DNA content, and the y axis refers to EdU incorporation. Overlay plots show colors in different cell cycle phases. (G1-blue; S-green; G2/M-orange) (g) Protein expression profile of mAID-ORC2^gr^, ORC2, ORC1, ORC3, CDC6, ATM, p-ATM(S1981), CHK1, p-CHK1(S345), p-CHK2(T68), and p-gH2AX(S139) in 4 cell lines after dox and auxin treatment for 0, 4, 24, and 50 hr. Cells were pretreated with doxycycline for 24 hr. before auxin treatment. Immunoblots for each protein were developed on the same film at the same time for comparison between four cell lines. (h) Immunofluorescent staining of p-ATM(S1981) in four cell lines with or without dox+auxin treatment. (i) Immunofluorescent staining of p-ATM(S1981) in four cell lines with or without dox+auxin treatment. For (h) and (i), dox+auxin treated cells were pre-treated with doxycycline for 24 hr. before adding auxin for 48 hours. Scale bar indicated 4 μM.

We focused on analysis of the effects of auxin-induced ORC2 depletion in the H-2, H-4, and H-5 cell lines because they have rapid depletion of mAID-ORC2^gr^ after auxin treatment. We excluded off target effects by confirming the ORC2-H-2 cloned cell line was resistant to both ORC2-1 and ORC2-2 sgRNAs, but not to the ORC2-3 and ORC2-4 sgRNAs (Figure 3—Figure supplement 1b). Compared with the parental TO-HCT116 cell line, the human diploid cell RPE-1 had about 50% of the amount of ORC2, while IMR-90 cells had about 15% (Figure 3a). The relative levels of ORC3 reflect the levels of ORC2 since they are known to form an intertwined complex throughout the cell cycle (Dhar et al., 2001; Jaremko et al., 2020; Vashee et al., 2001). ORC2_H-2, H-4, and H-5 had no endogenous ORC2 detected (note, a nonspecific smaller band was detected in between the two endogenous ORC2 proteins) (Figure 3a). In addition,

ORC2_H-2 cells expressed mAID-ORC2^gr^ at about 5% compared to endogenous ORC2 level in TO-HCT116, and H-4 and H-5 cells expressed about 10% of the endogenous ORC2 level. It is known that cancer cells can proliferate normally with 10% of the levels of ORC2 (Dhar et al., 2001).

Next, we compared the proliferation rates in these cell lines. In normal media conditions the ORC2_H-4, H-4, and H-5 cells grew only slightly slower than the parental TO-HCT116 cells (Figure 3b). When doxycycline only was added to induce the OsTIR protein expression, the proliferation rate of all cell lines decreased, possibly due to some toxicity of doxycycline or the expression of the OsTIR1 protein itself (Figure 3c). Importantly, the ORC2_H-4, H-4, and H-5 cells proliferated like the parental TO-HCT116 cells. Moreover, auxin alone did not affect the proliferation rate of wild type TO-HCT116, H-4, and H-5 cells, but it reduced the proliferation rate of H-2 cells substantially (Figure 3d). This phenotype was probably due to the leaky expression of Tet-OsTIR1 in the ORC2_H-2 cells. Importantly, when both doxycycline and auxin were added, all three ORC2 KO cell lines stopped proliferating, whereas the parental TO- HCT116 cells continued to proliferate (Figure 3e).

Concomitant with the lack of cell proliferation, the cell cycle profile changed after mAID-ORC2^gr^ was depleted in these cells. Cells were treated with doxycycline and auxin to deplete mAID- ORC2^gr^ for 0 hr., 4 hr., 24 hr., and 50 hr. at most. At the 50 hr. time point, all three ORC2 KO cell lines had less cells progressing from G1 into S phase, and more cells accumulated at the end of S phase or at the G2/M phase (Figure 3f, Figure 3—Figure supplement 2). Cells with a 4C DNA content (late S/G2/M phase) continued to incorporate EdU suggesting that DNA replication was not complete, even though the bulk of the genome was duplicated. This phenotype was consistent with previous observations that cells treated with *ORC2* siRNA arrested in interphase (70%) or as rounded, mitotic-like cells (30%) (Prasanth et al., 2004).

To analyze whether the cell cycle arrest was due to checkpoint activation in response to DNA damage, cell extracts were prepared from doxycycline and auxin treated cells and proteins detected by immunoblotting with various DNA damage markers. CHK1 is essential for the DNA damage response and the G2/M checkpoint arrest and is primarily phosphorylated by ATR, while phosphorylation by ATM has also been reported (Gatei et al., 2003; Goto et al., 2019; Jackson et al., 2000; Liu et al., 2000; Wilsker et al., 2008). ORC3 and mAID-ORC2^gr^ proteins in ORC2_H-2, H-4, and H-5 cell lines were depleted after 4 hours of auxin treatment, while ORC1 and CDC6 levels remained unchanged (Figure 3g). Phosphorylation of ATM(S1981), ATR(T1989), and CHK1(S345) was detected in H-2, H-4, and H-5 after 50 hr. of auxin treatment, but not in the parental TO-HCT116 cells (Figure 3g, Figure 3—Figure supplement 3a). Higher levels of P-gH2AX(S139) in H-2, H-4, and H-5 cells were detected even when no auxin was added (Figure 3g). This showed that although cells can divide with only 5-10 % of ORC2, a certain degree of DNA damage existed. In this experiment, the level of phosphorylated-CHK2(T68) showed no difference between control and mutant cells, but did increase along with mAID-ORC2^gr^ depletion. Supporting results were also found using immunofluorescent staining of individual cells. When doxycycline and auxin were added for 48 hours, substantially more ATM(S1981) and CHK1(S345) phosphorylation were detected in all three ORC2 KO cell lines (Figure 3h-i). In the absence of doxycycline and auxin, the P- gH2AX(S139) signal was more abundant in ORC2_H-2, H-4, and H-5 cells than in wild type (Figure 3—Figure supplement 3b). To conclude, insufficient ORC2 protein in cells resulted in abnormal DNA replication and DNA damage, and in response to DNA damage, CHK1 was activated and cells arrested in G2 phase.

### Loss of ORC2 results in heterochromatin decompaction and abnormal nuclear morphology

The ORC2 depleted ORC2_H-2 and ORC2_H-5 cells had twice the nuclear volume following treatment with doxycycline and auxin for 48 hr. (Figure 4a-b) compared to cells without treatment. The volume of the nucleus was greater than the volume of the largest parental cells, and thus was not due to their arrest with a 4C DNA content. This phenotype could be due to cells arrest in G2 phase with heterochromatin decompaction, since ORC2 depletion using siRNA decondenses centromere associated a-satellite DNA (Prasanth et al., 2010). During interphase, ORC2 and ORC3 localize to the heterochromatin foci and interact with heterochromatin protein 1a (HP1a) through ORC3 (Prasanth et al., 2004). To detect heterochromatin decompaction, immunofluorescent staining of the centromeric protein C (CENP-C) was performed. In TO-HCT116 cells, CENP-C staining showed multiple, compact foci, but in the doxycycline and auxin treated cells that were dependent on mAID-ORC2^gr^, CENP-C foci were larger and more prominent (Figure 4c). These observations were consistent with heterochromatin decompaction in ORC2 depleted cells.

**Figure 4.**
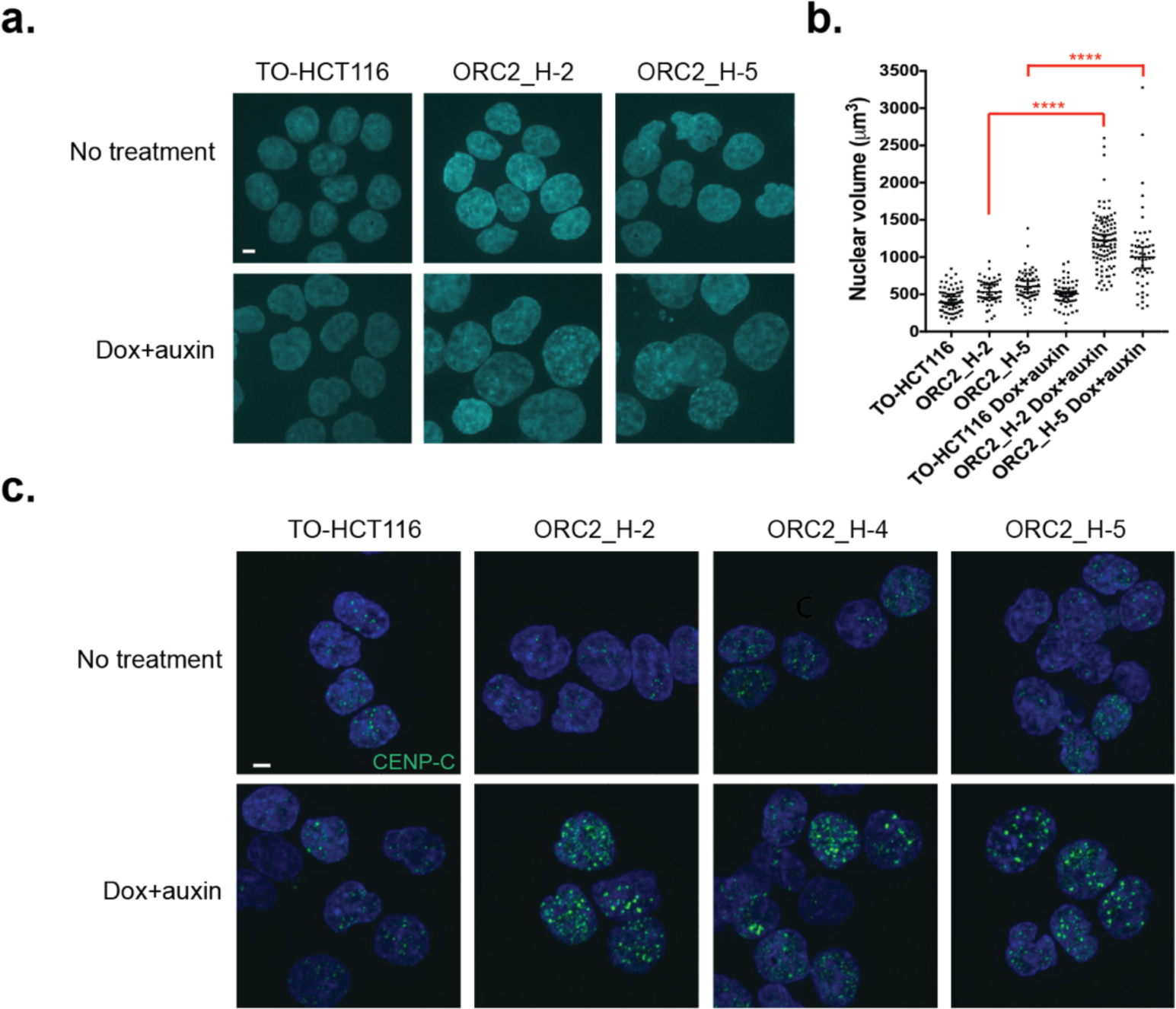
Auxin-treated ORC2_H2, H-4, and H-5 cells had abnormal nuclear phenotypes. (a) Nuclear morphology of TO-HCT116, ORC2_H-2 and H-5 cells after 48 hr. auxin treatment. Scale bar indicated 4 μM. (b) Scatter plot illustrating the nuclear volume after 48 hr. auxin treatment. Untreated: TO-HCT116, n=77; ORC2_H-2, n=52; ORC2_H-5, n=63. Dox and auxin treated: TO- HCT116, n=66; ORC2_H-2, n=110; ORC2_H-5, n=54. Error bars, medium ± 95% CI. Nuclear volume decreased significantly in both dox and auxin treated ORC2_H-2 and H-5 cells. (Student t-test, **** p<0.0001) (c) Immunofluorescent staining of CENP-C after mAID-ORC2^gr^ depletion for 50 hrs. Scale bar indicated 4 μM.

### ORC2 is essential for initiation of DNA replication

When cells were treated with siRNA against *ORC2* for 72 hours, 30% of the cells arrested in a mitosis-like state (Prasanth et al., 2004). This observation led to the conclusion that ORC2 is not only required for the initiation of DNA replication, but also during mitosis. To examine the role of ORC2 in G1 and mitosis following acute depletion, TO-HCT116 and the ORC2_H-2 cells were synchronized at the G1/S phase boundary with a 2C DNA content with a double thymidine block, with doxycycline added during the second thymidine block. When indicated, auxin was added 4.5 hours before the release from the second thymidine block and the cells followed for progression through two cell division cycles (Figure 5a). Cells were harvested at several timepoints after release and pulse-labeled with EdU for two hours. During the first cell cycle following release into S phase, no obvious change in DNA content and EdU labeling was observed, whether doxycycline and auxin was added or not (Figure 5b-c; Figure 5—Figure supplement 1). During the second cell cycle, however, doxycycline and auxin-treated TO- HCT116 cells progressed through S phase only slightly slower than the untreated cells. In contrast, serious cell cycle defects were observed between the ORC2_H-2 auxin-treated or untreated cells, starting with the second cell cycle (Figure 5b-c). First, auxin treated ORC2_H-2 cells exhibited a very slow S phase, indicating that cells were struggling to correctly replicate the DNA. Second, cells arrested with a 4C DNA content, which could be in the stage of late S or G2/M phase. Thirdly, after 48 hours of releasing from the double thymidine block, some cells were arrested in G1 phase and couldn’t enter S phase. Since auxin was added 4.5 hours before second thymidine release and it required about 4 hours to knockdown mAID-ORC2^gr^, those phenotypes were observed only during the second cell cycle, suggesting that, for the first cell cycle, the pre-RC was already formed and the cells were primed to replicate the complete genome. ORC2 depleted cells then either arrested during G1 or went through an incomplete S phase and arrested at the G2/M phase and did not progress further. This experiment indicated ORC2 primarily functions in establishing DNA replication initiation, as expected, but based on the results so far, we could not conclude a role during mitosis because most ORC2 depleted cells with a 4C DNA content continued to replicate DNA.

**Figure 5.**
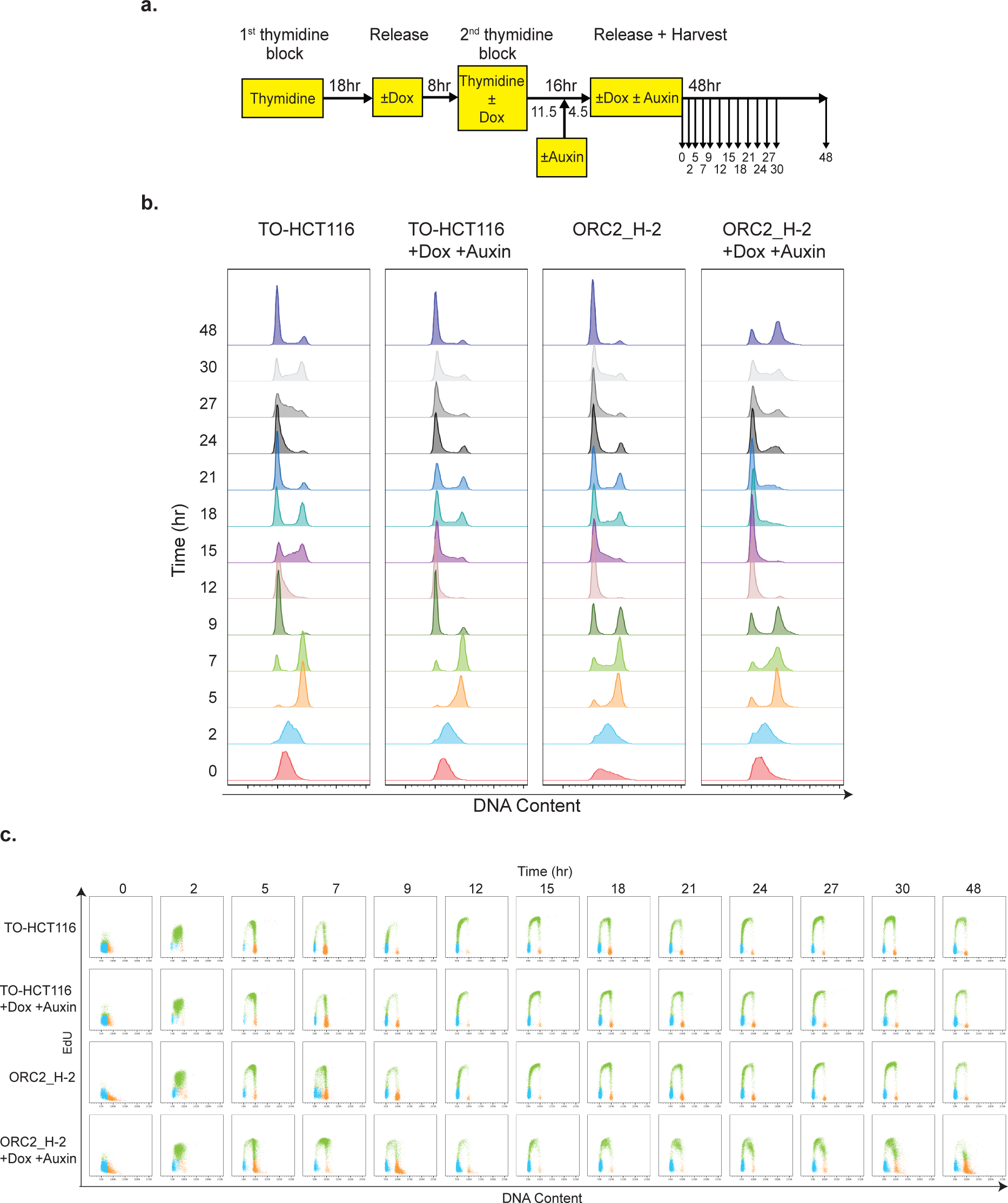
ORC2_H-2 cells show abnormal cell cycle progression after mAID-ORC2^gr^ depletion. (a) Experimental scheme of TO-HCT116 and ORC2_H-2 cells synchronization by a double thymidine block. (b) Flow cytometry analysis of FxCycle™ Violet stained cells (singlets) released from double thymidine block in indicated treatment. (c) Cell cycle analysis of TO- HCT116 and ORC2_H-2 cell lines released from double thymidine block in indicated treatment. Cells were pulse labeled with 10μM EdU for 2 hours before harvesting at reported time points. The x axis indicates DNA content, and the y axis refers to EdU incorporation. Overlay plots show colors in different cell cycle phases. (G1-blue; S-green; G2/M-orange)

### An MCM complex loading and pre-RC assembly defect in ORC2 depleted cells

The auxin-treated, mAID-ORC2^gr^ depleted cells could not replicate normally, possibly due to insufficient ORC to form the pre-RC. To test this hypothesis, the DNA-loaded MCM2-7 was measured by extracting the asynchronous cells with detergent and examining the chromatin bound MCM2 in relation to cell cycle stage using flow cytometry, as described previously (Matson et al., 2017). Asynchronous TO-HCT116, ORC2_H-2, and H-5 cells with or without doxycycline and auxin treatment were pulse-labeled with EdU, harvested and stained with anti- MCM2 antibody and DNA dye. The results showed that in normal media without detergent extraction, nearly 100% of the cells were positive for MCM2 in all three cell lines (Figure 6a; Figure 6 –Supplement 1). When extracted, about 78 % of TO-HCT116, 65.2 % of ORC2_H-2, and 76.9 % of ORC2_H-5 cells were positive for MCM2. When cells were treated with both doxycycline and auxin for 20 hours, 84.5 % of TO-HCT116 cells had DNA-loaded MCM2, while only 4.42% of H-2 and 26.2% of H-5 cells did (Figure 6b; Figure 6—Figure supplement 1). The different degrees of reduced MCM2 between H-2 and H-5 was expected because the level of mAID-ORC2^gr^ in H-2 was only half the amount compared to H-5 cells, and auxin knockdown was not 100% efficient. Of particular importance was the lack of MCM2 on chromatin in the G1 phase cells when mAID-ORC2^gr^ was depleted. In conclusion, low number of pre-RC formation on DNA origins results in cells arresting in G1. Those cells that still initiated DNA replication have a very slow S phase and arrested at G2/M with DNA damage and checkpoint activation.

**Figure 6.**
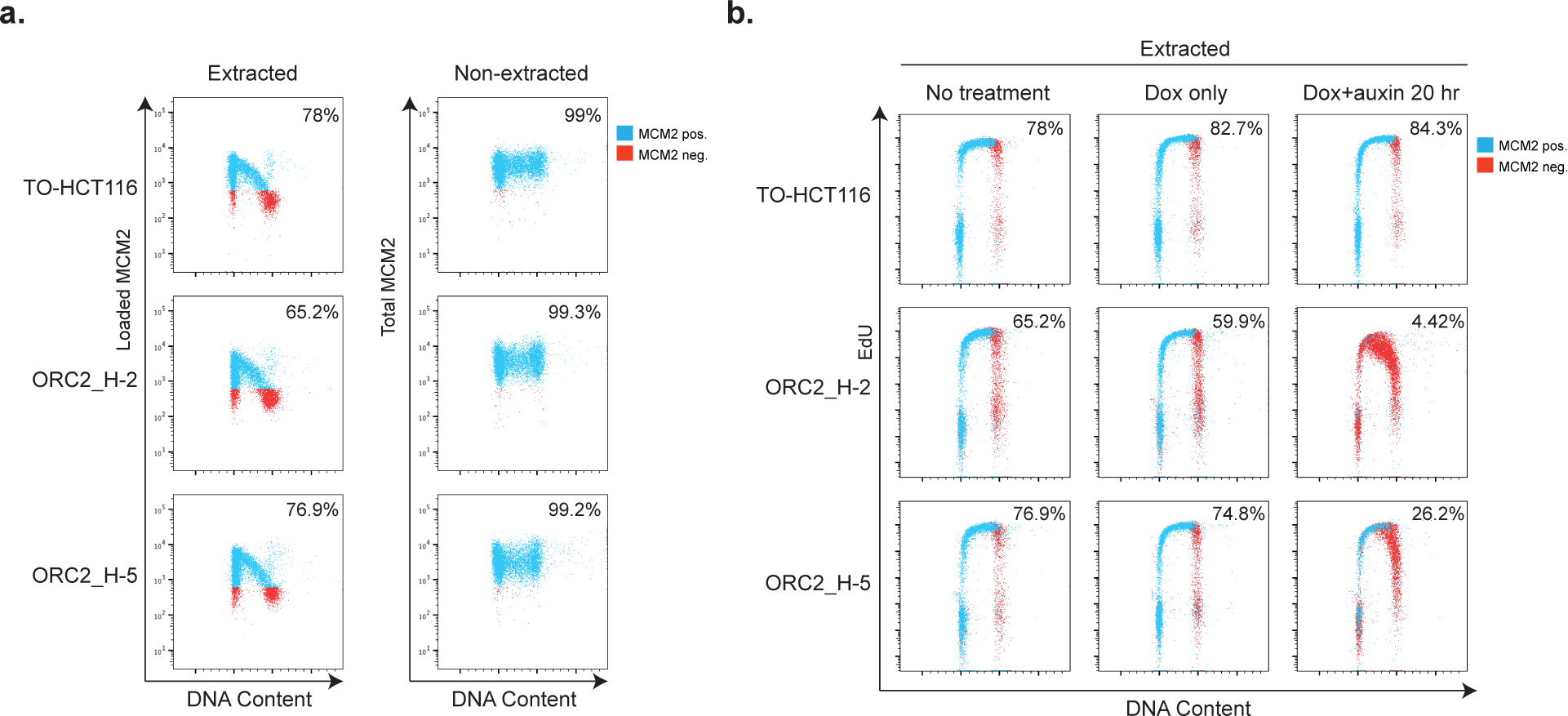
Depletion of mAID-ORC2^gr^ in ORC2_H-2 and H-5 cells result in decreased DNA-loaded MCM. (a) Flow cytometry analysis of DNA-bound MCM2 and total MCM2 in asynchronous cells. Extracted: Cells were treated with nonionic detergent to wash off unbound MCM2 before fixation, and then stained with anti-MCM2 antibody and Alexa Fluor 647- conjugated secondary antibody. Non-extracted: Cells were fixed right after harvest and detected for MCM2 with the same procedure. Blue cells are MCM2 positive, and red cells are MCM2 negative. The x axis indicates DNA content, and the y axis refers to MCM2 level. (b) Flow cytometry measuring DNA content, EdU incorporation, and DNA-bound MCM2 in asynchronous cells in different condition. Cells were pulse labeled with 10 μM EdU for 2 hours before harvesting. The x axis indicates DNA content, and the y axis refers to EdU incorporation. MCM2 positive and negative cells are shown as blue and red cells. Numbers at the upper right corner indicates percentage of MCM2 positive population.

### ORC2 depletion in cells leads to aberrant mitosis

In order to know if the mAID-ORC2^gr^ depleted cells entered mitosis, we evaluated the mitotic index by staining cells with anti-pH3S10 and followed by flow cytometry analysis. In a normal asynchronous cell population, about 4.53 ± 0.59 % and 1.57 ± 0.33 % were pH3S10-positive in TO-HCT116 and ORC2_H-2 cells respectively, while 31.4 ± 2.88 % of TO-HCT116 cells and 15.6 ± 1.25 % of ORC2_H-2 cells were at G2/M (Figure 7a; Figure 7—Figure supplement 1). When only doxycycline was added, there was no significant change. When treated with doxycycline and auxin for 28 hours, the pH3S10-positive cell population percentage was about 2.39± 0.26 in TO-HCT116 and only 0.79 ± 0.09 in ORC2_H-2, while 17.23 ± 0.78 % of TO- HCT116 cells and 36.7 ± 1.61 % of ORC2_H-2 cells were at G2/M. After 50 hr. dox and auxin treatment, the pH3S10-positive cell population percentage was 3.95 ± 0.16 in TO-HCT116 and only 0.96 ± 0.15 in ORC2_H-2, while 20.77 ± 1.76 % of TO-HCT116 cells and 79.57 ± 1.2 % of ORC2_H-2 cells were at G2/M phase. In normal media conditions, TO-HCT116 already had 2.9 times as many mitotic cells as ORC2_H-2. When treated with doxycycline and auxin, although the G2/M population increased 2.3- fold and 5- fold at 28 hr. and 50 hr., respectively, the number of mitotic cells in ORC2_H-2 was reduced 50 - 80 % compared to the non-treated H-2 cells. This showed that most ORC2_H-2 cells accumulated at the 4C DNA peak after ORC2 depletion were in the G2 stage and most cells did not enter into mitosis.

**Figure 7.**
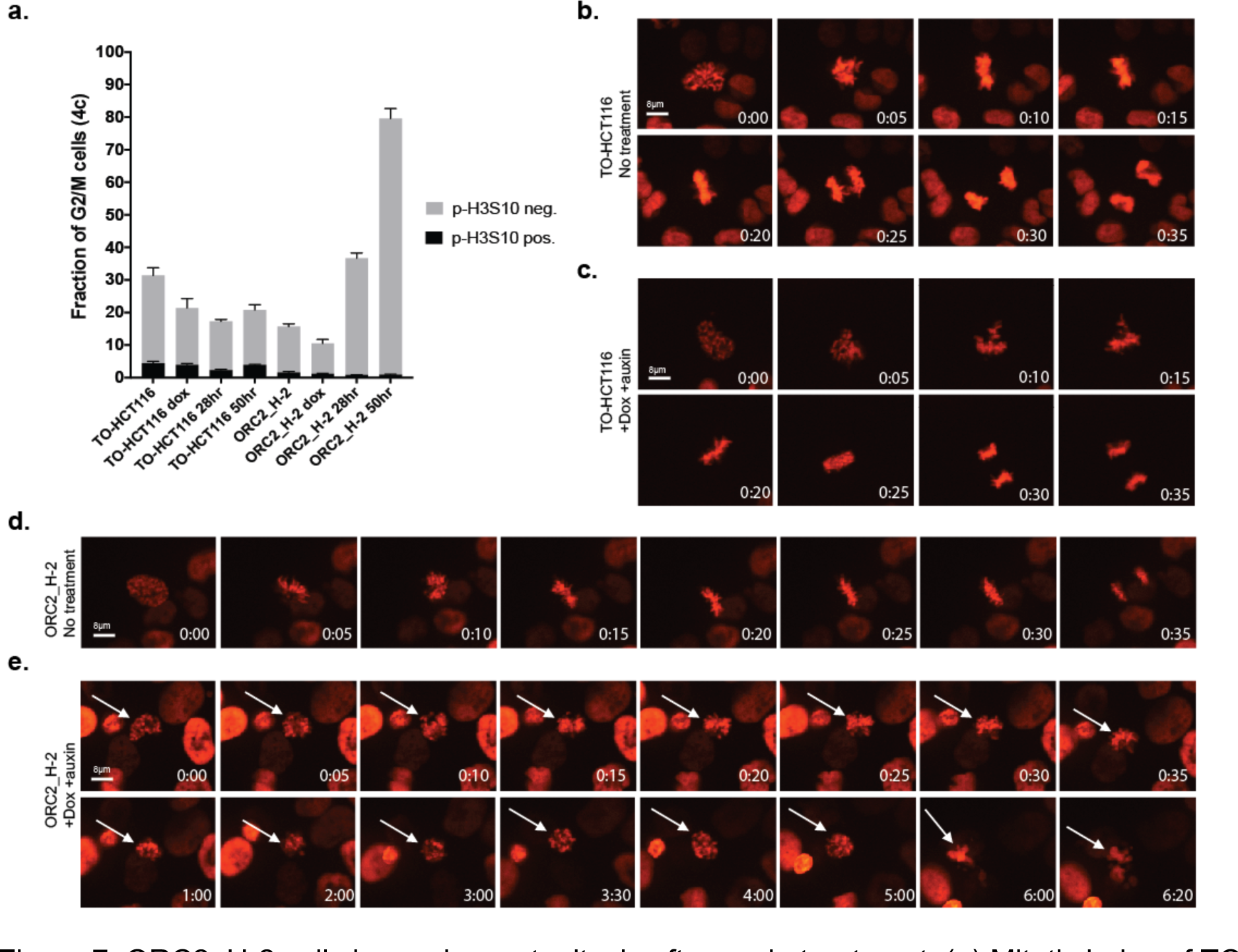
ORC2_H-2 cells have aberrant mitosis after auxin treatment. (a) Mitotic index of TO- HCT116 and ORC2_H-2 G2/M cells with or without auxin. 0.75 µg/ml Doxycycline were added for 24 hr. before auxin treatment. Cells were harvest after 0, 28, or 50 hr. of auxin treatment followed by staining with anti-pH3S10 antibody for mitotic cells and FxCycle™ Violet for DNA content. The x axis refers to cell line and different conditions, including no treatment, doxycycline only, dox+auxin for 28 hr, and dox+auxin for 50 hr. The y axis is the fraction of 4c G2/M cells. Cell population positive or negative for p-H3S10 were shown as black or grey color. N=3 (biological repeats). (b-e) Time lapse imaging of TO-HCT116 and ORC2_H-2 cell lines. Cells were first arrested with single thymidine block (± dox) for 24 hr. and then released into desired medium. Time shown in lower left corner indicates time (hour : minute) since early prophase. (b) Images of TO-HCT116 cells without auxin were taken at 36 hr. 33 min. after released from thymidine block. (c) Auxin treated TO-HCT116 cells were taken at 32 hr. 58 min. after released from thymidine block. (d) ORC2_H-2 cells were taken at 41 hr. 38 min. after released from thymidine block. (e) Dox and auxin treated ORC2_H-2 cell were taken at 33 hr. 44 min. after released from thymidine block. White arrows in (e) pointed to the same cell. Scale bar indicated 8 μM.

Nevertheless, mitosis still occurred at a very low frequency. To observe how mitosis progression was affected after ORC2 depletion, we constitutively-expressed H2B-mCherry in TO-HCT116 and ORC2_H-2 cells via lentiviral transduction and performed time lapse fluorescent imaging of the mitotic chromosomes. Cells were synchronized using a single thymidine block and auxin was added or omitted 2 hours before releasing into fresh media with or without doxycycline and auxin. As expected, the first cell cycle after releasing from the thymidine block in both cell lines with or without doxycycline and auxin was normal and cells that progressed into mitosis and into the second cell cycle. During the second cell cycle, in the absence or presence of doxycycline and auxin, it took TO-HCT116 cells about 35 min. to progress from prophase to chromosome segregation (Figure 7b and c). The ORC2_H-2 cells in the absence of doxycycline and auxin also progressed thought mitosis like the parental cells (Figure 7d). In stark contrast, the ORC2_H-2 cells in the presence of doxycycline and auxin condensed chromatin and attempted to congress chromosomes at the metaphase plate but never achieved a correct metaphase alignment of chromosomes even after seven hours (Figure 7e). Those few cells that did attempt anaphase had abnormal chromosome segregation, producing lagging chromosomes, micronuclei and eventually apoptosis.

### Characterization of previously published *ORC1^-/-^* and *ORC2^-/-^* cell lines

The results so far confirm previous observations that ORC is essential for cell proliferation in human cells, but there remains the curious case of the viable knockout of *ORC1* and *ORC2* genes in p53^-/-^ HCT116 cells which needs to be explained (Shibata et al., 2016). We obtained two of the *ORC1^-/-^* and *ORC2^-/-^* cell lines used in that study and performed several experiments. We first tested whether using CRISPR/Cas9 to target *ORC2* with sgRNAs in the *ORC2^-/-^* cell line, whether negative selection occurred like that in the wild type HCT116 cells as shown in (Figure 2e). The sgRNA target GFP competition assay showed that in both parental HCT116 p53^-/-^ and *ORC2^-/-^* cell lines, cells targeted with *ORC2* sgRNA were outgrown by sgRNA-negative cells (Figure 8a-b). More importantly, both cell lines expressing mAID-ORC2^gr^ rescued or partially rescued the sgRNA targeting effect with a mAID-ORC2^gr^ that was resistant to the ORC2-1 and ORC2-2 sgRNAs, showing that the depletion was not due to an off-target effect of the *ORC2* sgRNAs (Figure 8c-d). This assay suggested that there is some form of functional ORC2 in the *ORC2^-/-^* cells that could be targeted by the tested sgRNAs. In addition, an immunoblot of the cell lysates showed a reduced level of ORC3 in the *ORC2^-/-^* cells, and since ORC2 and ORC3 form stable heterodimers in cells, this result again indicated that some form of ORC2 was expressed in cells, albeit at a lower level (Figure 8—Figure supplement 1a). When immunoprecipitated with an antibody against ORC3, we detected ORC3 and a putative truncated form of ORC2 which was seen only in *ORC2^-/-^* cells (Figure 8—Figure supplement 1b). Next, we designed primer pairs that span the exon junction for every exon in *ORC2* and performed quantitative RT-PCR (qPCR) to determine the nature of the *ORC2* transcripts in the *ORC2^-/-^* cells (Figure 8e). The fold change (FC) indicated that in *ORC2^-/-^* cells, about 60% of the mRNAs had exon 7 skipped, whereas other exons remained the same (Figure 8—Figure supplement 2).

**Figure 8.**
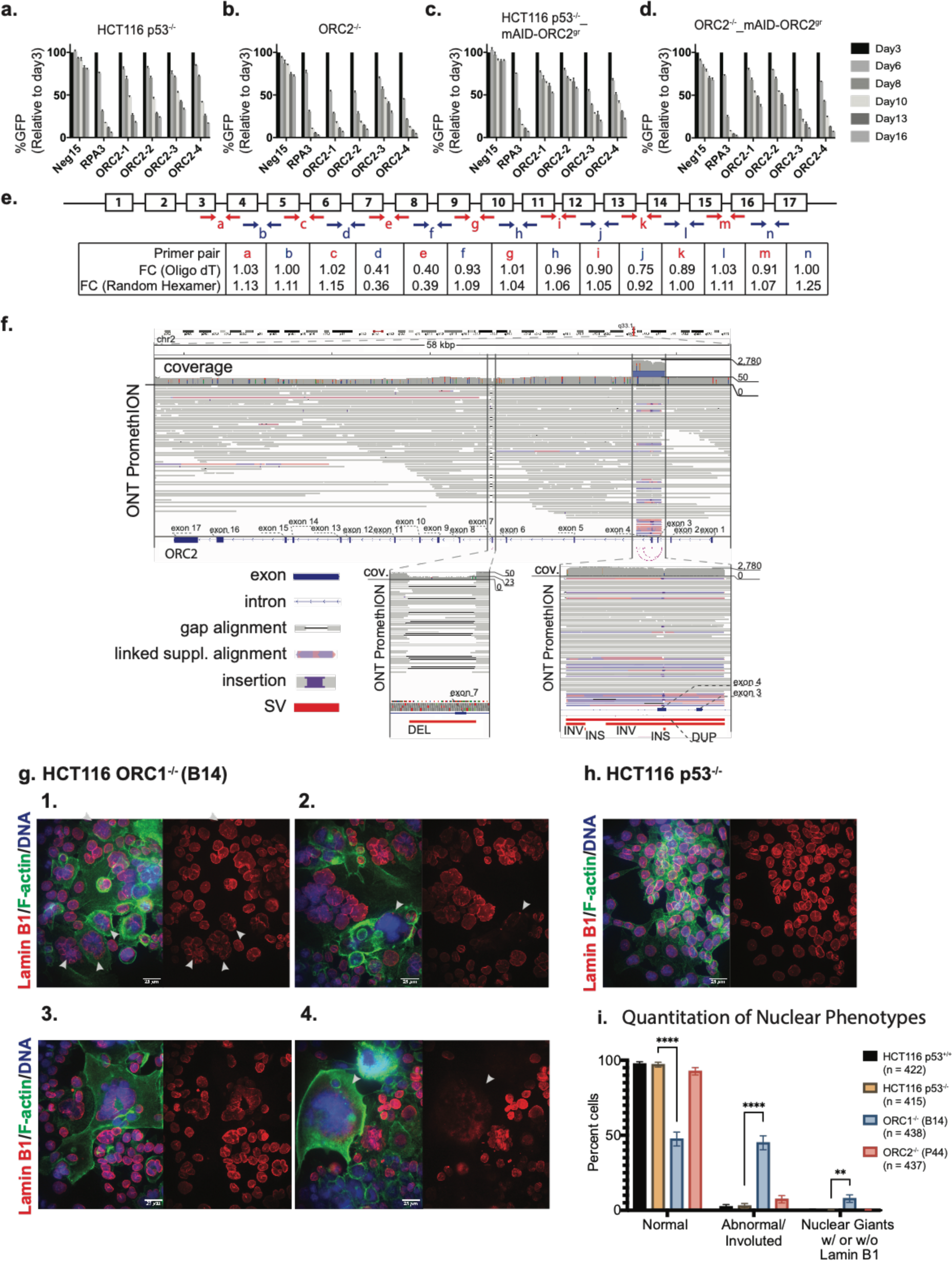
Characterization of previously published *ORC1^-/-^* and *ORC2^-/-^* cell lines. (a-d) Negative-selection time course assay that plots the percentage of GFP^+^ cells over time following transduction with the indicated sgRNAs with Cas9. Experiments were performed in HCT116 *p53^-/-^*, *ORC2^-/^*^-^, HCT116 *p53^-/-^*_mAID-ORC2^gr^, and *ORC2^-/-^*_mAID-ORC2^gr^ cell lines. The GFP positive percentage was normalized to the Day3 measurement. *n* = 3. Error bars, mean ± SD. (e) Calculated fold change (FC) for each primer pairs in ORC2^-/-^ cells compared to HCT116 p53^-/-^ cells. The red and blue arrows indicate each the primer pair. Two kinds of primers, Oligo dT and Random Hexamer, were used in the reverse transcription step. Bar diagram view is shown in Figure 8—Figure supplement 1. (f) Structural Variations (SVs) in *ORC2* gene. Top panel shows the alignment overview of the ONT Promethion long reads over the *ORC2* gene region with coverage track showing an average coverage of ∼50x for the majority of the gene sequence, with exceptions in regions around exon 3, 4, and 7. Zoomed-in region with heterozygous deletion spanning the exon 7 of the *ORC2* gene, with supporting gap aligned ONT reads and coverage track with a characteristic “dip” over the deleted segment. Zoomed-in rearranged region around exons 3 and 4 of the *ORC2* gene, with linked supplement read alignments pileup over the affected area supporting the spanning duplication SV, with 2 more nested inversions and 2 more nested insertion highlighted in the SV track. (g) 1-4: Immunofluorescence of HCT116 *ORC1^-/-^* (B14) cell line stained with Anti-Lamin B1 antibody (Red), Phalloidin (F-actin) (Green), Hoechst Dye (Blue). Images show either merge of all three channels or Lamin-B1 staining of the nuclei. White arrows indicate abnormal and involuted nuclei in image 1. White arrows also show extremely large (nuclear giants) that have lost nuclear membrane integrity (2,4). (h) parental cell line for the *ORC1^-/-^* line presented as control for quantitative and qualitative comparison. More fields of control cells HCT116 *p53* WT and *p53* null background, HCT116 *ORC1^-/-^* and HCT116 *ORC2^-/-^* cell lines are shown in Figure 8—Figure supplement 3 and 4. Scale bar is 25 µm (i) Quantitation of abnormal nuclei between cell lines. Significance calculated using two-way ANOVA for multiple comparisons keeping HCT116 p53^-/-^ as control. **** p < 0.0001, ** p = 0.0028.

With the idea that genomic instability caused by the absence of or mutation within either *ORC1* or *ORC2* in these cell lines might give rise to copy number variations, we performed SMASH (Wang et al., 2016) analysis on the two parental HCT116 with the p53 WT and p53 null background as well as the Shibata et al. (2016) *ORC1* and *ORC2* deficient lines. Both the parental cell HCT116 cell lines showed very similar chromosome copy number, while both *ORC1* deficient and *ORC2* deficient cell lines had additional CNVs in chromosomes unrelated to those harboring either *ORC1* or *ORC2* (Figure 8—Figure supplement 3). The significance of these specific loci which showed alterations in copy number when either *ORC1* or *ORC2* was deleted remains to be seen. However, it was in this analysis that we also noticed that part of the *ORC2* gene locus was hugely amplified (Figure 8—Figure supplement 3 solid arrow). To study in detail the *ORC2* gene region on chromosome 2 in the putative ORC2^-/-^ cells, we performed long-read Nanopore sequencing analysis that showed the *ORC2* gene in *ORC2^-/-^* cells was highly rearranged and heterogenous (Figure 8f). Aside from the aforementioned heterozygous deletion of the 7^th^ exon, the region near the 3^rd^ and 4^th^ exons also exhibited overwhelming amplification and structural rearrangement signatures (Figure 8f). Thus, the *ORC2* gene is present but has heterogenous expression of exon 7 and is sensitive to sgRNAs that target *ORC2*. We conclude that the cell line is not a true *ORC2* knockout.

With regard to the HCT116 *ORC1^-/-^* (B14) cell line, we confirmed that they lacked ORC1 protein using multiple antibodies and confirmed that they duplicated at a much slower rate than the parental line, as previously reported (Shibata et al., 2016). Furthermore, we were unable to passage these cells for many generations (by 30 generations they stopped proliferating). We also compared HCT116 *ORC1^-/-^* (B14) and HCT116 *ORC2^-/-^* (P44) cell lines with the parental lines, both either p53 WT or null background by confocal microscopy [Figure 8g (1-4), 8h; Figure 8—Figure supplement 4a-e, Figure supplement 5a-c]. There was a myriad of nuclear morphology defects in the *ORC1* deficient cell line. When the nuclei were stained with Hoechst dye, up to 10 percent contained abnormally large nuclei or sometimes what seemed to be multiple nuclei aggregated together in single cell, while many other of the cells appeared normal. When probed further by staining for F-actin and Lamin B1 for overall cellular and morphology and nuclear membrane integrity respectively, we saw that despite the staining for DNA content looking normal, up to 50 percent of the *ORC1^-/-^* cells showed highly abnormal, involuted nuclear membranes (Figure 8i). In addition, most of the gigantic nuclei seemed to have lost the nuclear membrane, while those cells that had Lamin B1 staining displayed abnormal nuclear membrane integrity.

The chromatin organization in the *ORC1* deficient cells was observed by transmission electron microscopy (TEM) and revealed a huge difference in cell size and nuclear structure between the wild type HCT116 *p53^+/+^* and *ORC1^-/-^* (Figure 8—Figure supplement 5). About 35 % of *ORC1^-/-^* cells were grossly larger than wild type cells. Those multinucleate/polyploid giant cells were full of membrane invagination, vacuoles and apoptosis. Most likely they were formed due to extensive DNA damage and nuclear structural defects and underwent a different type of cell division called neosis, in which intracellular cytokinesis occurs and some mononuclear cells are produced from nuclear budding or asymmetric cell division (Sundaram et al., 2004). All those phenotypes pointed to the fact that although *ORC1^-/-^* cells do not survive in culture in the long term and even when they were slowly proliferating, they were grossly abnormal. It may well be the case that the p53^-/-^ status was required for these cells to be produced in the first place.

## Discussion

The *ORC2^-/-^* cell line believed to be a complete knockout via the use of 3 sgRNAs, one targeting the exon 4, and the others targeting the 6^th^ and 7^th^ introns retained a truncated form of ORC2 that could interact with ORC3 and was expressed from a rearranged gene. These cells were still susceptible to ORC2 knockdown using four sgRNAs selected from our CRISPR screens and also partially rescued the phenotype with of two sgRNAs using a CRISPR-sgRNA resistant mAID- ORC2^gr^. Similar to what we found for *ORC2^-/-^* cells, CRISPR-induced frameshifts in cells often generate truncated proteins that, although they may not be recognized by western blot, still preserve whole or partial protein function (Smits et al., 2019). Based on these observations with ORC2 and the results with *ORC1* deficient cells, that despite the over-production of CDC6, were unable to proliferate for many generations and produced abnormally structured cells, as well as data analyzed by tiling-sgRNA CRISPR screens, we conclude that ORC is essential in human cells. This conclusion is consistent with existing literature (Hemerly et al., 2009; McKinley and Cheeseman, 2017; Ohta et al., 2003; Prasanth et al., 2010, 2004, 2002) and is not surprising since ORC has multiple functions in human cells and ORC is essential in all other eukaryotic cells examined.

The pooled CRISPR/Cas9 domain-focused screen has become a common and powerful tool for uncovering genes that are essential for cell proliferation, cell survival, and for identification of essential functional domains in proteins (Adelmann et al., 2018; Park et al., 2016; Shi et al., 2015; So et al., 2019). However, if the screens use only a handful of guides targeting annotated essential regions, it may still result in data which may or may not score a gene as essential. Tiling-sgRNA CRISPR-Cas9 screens on the other hand test ‘functional’ or ‘essential’ domains in a more unbiased way. Using this approach targeting sgRNAs across entire open reading frames of ORC1-6 and CDC6 enabled us to classify them as essential, and correlate functional domains within these proteins. The combined results also confirmed that all ORC1-6 and CDC6 proteins were essential in cancer cells as well as a human diploid cell line, including ORC2 that was characterized in the DepMap portal (https://depmap.org/portal/) as non-essential based on multiple shRNA and whole genome CRIPSR-Cas-9 screens in multiple cells. We were able to identify many sgRNAs that targeted *ORC2* in the tiling-sgRNA CRISPR screen and the two chosen cloned sgRNAs that killed cells were successfully complemented using a *mAID-ORC2^gr^* transgene, demonstrating specificity of the knockdowns. Thus, large scale experiments, especially negative results, should be interpreted with caution, such that the essential nature of a gene should be examined in depth as we have done here.

The known functional domains in ORC1, including the BAH, AAA+ and WHD were identified using the open reading frame tiling CRISPR-Ca9 sgRNA screen, as well as other regions of ORC1, including the intrinsically disordered region (IDR; amino acids 180-480, Figure 1e) which we know binds Cyclins E and A-CDK2 and CDC6 (Hossain et al., 2019) as well as many other proteins we have identified and are characterizing in detail. The screen also identified an essential region of ORC1 in and around amino acid 750-790 (Figure 1a-b) which may represent the pericentrin- AKAP450 centrosomal targeting (PACT) domain that localizes ORC1 to centrosomes to regulate correctly centrosome and centriole copy number (Hemerly et al., 2009).

In ORC2, multiple, essential domains were identified, including the AAA+-like domain and the WHD. The WHD of human ORC2 controls access of human ORC to DNA by inserting itself into the DNA binding channel prior to activation of the protein by binding of ORC1 and subsequent binding of CDC6 (Hossain et al., 2019; Jaremko et al., 2020). The ORC2-carboxy terminus binds to ORC3 and ORC2 is also known to bind to PLK1, the mitotic protein kinase (Song et al., 2011). Interestingly, ORC2 also has an IDR (Figure 2—Figure supplement 1d; amino acids 30-230) and a gRNA tiling screen of this region shows limited essential amino acids, but a conserved region surrounding amino acid 190 is reproducibly essential in both HCT116 cancer cells and diploid (Figure 2a-b and Figure 2—Figure supplement 1a). The entire IDR, however, may contribute to DNA mediated ORC liquid phase transition (Parker et al., 2019).

The use of a mAID-ORC2^gr^ enabled rapid removal of ORC2 from cells and analysis of the resulting phenotypes. It was not surprising that ORC2 is essential for loading MCM2, and hence MCM2-7, to establish pre-RCs and origins of DNA replication across the genome. In the absence of ORC2, cells loaded little MCM2, most likely resulting in too few origins of replication and a consequent slow S phase and arrest with a near 4C DNA content and ongoing DNA synthesis. ORC2 depletion yielded other phenotypes, including large nuclei and a failure to execute mitosis. The large nuclei, also observed in the *ORC1^-/-^* cells, have large CENP-C foci, probably due to decompaction of the centromeric associated a-satellite DNA, as observed previously (Prasanth et al., 2010). We suggest a general role for ORC in nuclear organization and organizing chromatin domains in the nucleus, including heterochromatin. In yeast, ORC is essential for transcriptional silencing at the silent mating type heterochromatic loci *HMRa* and *HML*a loci and its function in replication are separable from that in silencing (Bell et al., 1993; DeBeer et al., 2003; Ehrenhofer-Murray et al., 1995). In *Drosophila*, ORC localizes and associates with heterochromatin protein HP1 during interphase and mitosis and heterozygous, recessive lethal mutations in *DmORC2* suppress position effect variegation (Huang et al., 1998; Pak et al., 1997). In human, ORC1 interacts with RB and SUV39H1, a histone methyltransferase that tri-methylates histone H3K9 which HP1 binds to repress E2F1-dependent CCNE1 transcription (Hossain and Stillman, 2016) . ORC1 and ORC3 (a tight ORC2 binding partner) directly interact with HP1, and depletion of ORC subunits disrupt localization of HP1 and the compaction of chromosome 9 a-satellite repeats DNA (Prasanth et al., 2010). The mechanism by which the nuclei become large as a result of ORC depletion is under further investigation.

A final phenotype we observed in the acute removal of ORC2 is that the cells that replicate DNA and enter into mitosis attempt chromosome congression at the metaphase plate, but never make it, even after 7 hours. Eventually the cells die of apoptosis. We had observed abnormal mitotic cells following long term (72 hr.) treatment of cells with shRNAs that targeted *ORC2* but it was not clear if this phonotype was due to incomplete DNA replication (Prasanth et al., 2004). However, in the current study, acute removal of ORC2 captured some cells with a clear defect in chromosome congression during mitosis. Moreover, both ORC2 and ORC3 localize to centromeres (Craig et al., 2003; Prasanth et al., 2004), suggesting that they play a role in spindle attachment or centromeric DNA organization, particularly the centromere associated satellite repeat sequences. We speculate that in ancestral species, ORC localized at origins of DNA replication and this ORC also functioned in organization of chromosomes and in chromosome segregation, but upon separation of DNA replication and chromosome segregation with the advent of mitosis, separate functions of ORC in DNA replication, chromatin or nuclear organization and chromosome segregation were retained, but executed at different times during the cell division cycle.

## Materials and methods

### Cell Culture

HCT116 (WT cell lines is *p53^+/+^*), U2OS, and RPE1 cell lines were obtained from the Cold Spring Harbor Laboratory and cultured in DMEM (Gibco) and supplemented with 10% Fetal bovine serum and 1 % Penicillin/Streptomycin. IMR-90 cell line is culture in EMEM with 10 % Fetal bovine serum and 1 % Penicillin/Streptomycin. Plat-E cells and HEK293T cells were cultured in DMEM supplemented with 10 % FBS and penicillin/streptomycin. Plat-E cells were used for retroviral production and HEK293T cells were used for lentiviral production. HCT116 (*p53^-/-^*), HCT116 *ORC1^-/-^* (*p53^-/-^* background, clone B14), HCT116 *ORC2^-/-^* (*p53^-/-^* background, clone P44) were a kind gift from Dr. Anindya Dutta (University of Virginia, Charlottesville, VA, USA). Tet-OsTIR1 HCT116 (TO-HCT116) cell line was a kind gift from Dr. Masato Kanemaki (National Institute of Genetics, Mishima, Japan). All gifted cell lines were cultured in McCoys 5A (Gibco) supplemented with 10 % fetal bovine serum and 1% Penicillin/Streptomycin. All cell lines were cultured at 37 °C with 5 % CO2. All of the cell lines used in this study were tested for mycoplasma and were negative.

### Tiling-sgRNA guide design

Every possible guide directly upstream of a sp-Cas9 canonical PAM (NGG) sequence in the 5’->3’ direction is extracted from the target exon sequences. Guides with the canonical PAM (NGG) are aligned to the hg38 genome using the BatMis exact k-mismatch aligner (Tennakoon et al., 2012). A maximum of three mismatches are considered for off-target evaluation. The resulting alignment file is parsed, and each off-target location is assessed a penalty according to the number of mismatches to the target sequence, the exact position of each mismatch in the guide, where the further the mismatch is from the PAM the higher the penalty, the proximity of the mismatches to each other; assigning higher penalties to mismatches that are further apart.

The resulting penalties from each assessed off-target site are then combined in to a single off-target score for each guide similar to (Hsu et al., 2013), with 1.00 as the maximum possible score for guides not having any off-target site with up to three mismatches. The final results include the guide sequence, the PAM, the number of off-target sites in the genome with 0, 1, 2 and 3 mismatches, the cut site location, the calculated off-target score, and any yes (Supplement Table 1_guides).

### Plasmid construction and sgRNA cloning

Clonal HCT116-Cas9 expressing cell lines were a gift from Dr. Chris Vakoc and RPE1-Cas9 expressing cell lines were derived from Dr. Jason Sheltzer (Cold Spring Harbor Laboratory, NY, USA). In this study, all the sgRNAs targeting genes of interest as well as controls were cloned into LRG2.1 (derived from U6-sgRNA-GFP, Addgene: 108098 - as described in ref (Tarumoto et al., 2018)). Single sgRNAs were cloned by annealing sense and anti-sense DNA oligos followed by T4 DNA ligation into a BsmB1digested LRG2.1 vector. To improve U6 promoter transcription efficiency, an additional 5’ G nucleotide was added to all sgRNA oligo designs that did not already start with a 5’ G. Sequences of all arrayed sgRNA libraries used in this study are provided in (Supplement Table1_guides)

For an unbiased tiling CRISPR domain screen, pooled sgRNA libraries were constructed. All possible sgRNAs (PAM - NGG) were designed across each exon of the 7 target genes. Targeting or positive/negative control sgRNAs were synthesized in duplicate or triplicate in a pooled format on an array platform (Twist Bioscience) and then PCR cloned into the Bsmb1- digested LRG2.1 vector using Gibson Assembly. To ensure the representation and identity of sgRNA in the pooled lentiviral libraries, a deep-sequencing analysis was performed on a MiSeq instrument (Illumina) and verified that 100 % of the designed sgRNAs were cloned in the LRG2.1 vector and that the abundance of >95 % of the sgRNA constructs was within 5-fold of the mean. While this was as a means to QC for ORC1-6 and CDC6 libraries, for time considerations, in case of MCM2-7, CDT1 and Control libraries, presence of all guides in the T=0 sampling during the pooled CRISPR screening (described later) served as a similar QC.

For CRISPR complementation assays, *ORC2* sgRNA resistant synonymous mutations were introduced to ORC2 by PCR mutagenesis using Phusion high fidelity DNA polymerase (NEB). Guide RNA resistant ORC2 (ORC2^gr^) was amplified by PCR and cloned into NheI-digested mAID-mCherry2-NeoR plasmid (mAID-mCherry2-NeoR, Addgene 72830) in order to add mAID degron sequence to the N-terminus. The mAID-ORC2^gr^ was then PCR and assembled into BglII/XhoI digested pMSCV-hygro retroviral vector (TaKaRa #634401). All cloning was done using In-Fusion cloning system (TaKaRa). In this experiment, sgRNAs targeting ORC2 and control sgRNAs were cloned into BsmB1digested LgCG_cc88 lentiviral vector by the same sgRNA cloning strategy described above.

For knocking out endogenous *ORC2* in TO-HCT116 cells, we used sgRNA_ORC2-1-epCas9-1.1-mCherry plasmid to perform CRISPR/Cas9 in the cells. Sequence of sgRNA_ORC2-1 was cloned into epCas9-1.1-mCherry plasmid which was a kind gift from Dr. David Spector (Cold Spring Harbor Laboratory, NY, USA). Single sgRNA were cloned by annealing sense and anti- sense DNA oligos followed by T4 DNA ligation into a BbsI -digested sgRNA_ sgRNA_ORC2-1- epCas9-1.1-mCherry vector.

To construct lentiviral vector that constitutively express H2B-mCherry in TO-HCT116 and ORC2_H-2 cells, H2B-mCherry sequence were PCR and cloned into BamHI/BspDI -digested pHAGE-CMV-MCS-IZsGreen vector which was a kind gift from Dr. Alea Mills (Cold Spring Harbor Laboratory, NY, USA).

### Viral Transductions

Lentivirus was produced in HEK293T cells by co-transfecting target plasmid and helper packaging plasmids psPAX2 and pVSVG with polyethylenimine (PEI 25000, cat#) transfection reagent. HEK293T cells were plated a day before to achieve 80 % - 90 % confluency on the day of transfection. Plasmids were mixed in the ratio of 1:1.5:2 of psPAX2, pVSVG and target plasmid DNA in OptiMEM (Cat#). 32 µl of 1 mg/mL PEI (calculated based on the final volume of transfection) was added, mixed and incubated, before addition to the cells. Cell culture medium was changed 7 h after transfection, then supernatant collected at 36 and 72 h following transfection was pooled. For the high throughput lentiviral screening, virus supernatant was concentrated with Lenti-X™ Concentrator (Takara, #631231) as per the manufacturer’s protocol.

Retrovirus was produced in Plat-E cells by co-transfecting target plasmid and packaging plasmids pCL-Eco and pVSVG in the ratio of 1.25:1:9 with PEI. Cell culture medium was changed 7 h after transfection, and the supernatant was collected at 36 hr. post-transfection.

For either lenti- or retroviral transductions, target cells were mixed with viral supernatant, supplemented with 8 µg/mL polybrene and centrifuged at 1700 rpm for 30 min. at room temperature. Fresh medium was replaced 24 h after transduction. Antibiotics (1 µg/mL puromycin; 10 µg/mL of blasticidin; 200 μg/ml of hygromycin) were added 72 h after infection when selection was required.

### Pooled sgRNA screening

CRISPR-based negative selection screenings using sgRNA libraries targeting proteins ORC1-6, CDC6 as well as positive and negative controls, were performed in stable Cas9 expressing HCT116 and RPE1 cell lines. The screens were performed as previously described (Lu et al., 2018; Miles et al., 2016; Shi et al., 2015) with a few optimizations for scale. Briefly, to ensure a single copy sgRNA transduction per cell, multiplicity of infection (MOI) was set to 0.3-0.35. To achieve the desired representation of each sgRNAs during the screen, the total number of cells infected was determined such that while maintaining the MOI at ∼0.3, the sgRNA positive cells were at least 2000 times the sgRNA number in the library. Cells were harvested at day 3 post-infection and served as the initial time-point (T=0) of the pooled sgRNA library, representing all guides transduced to begin with. Cells were cultured for 10 population doublings (T=10) and harvested as the final time point. All experiments were performed in triplicates. Genomic DNA was extracted using QIAamp DNA midi kit (QIAGEN) according to the manufacturer’s protocol.

Next Generation Sequencing library was constructed based on a newly developed protocol. To quantify the sgRNA abundance of initial and final time-points, the sgRNA cassette was PCR amplified from genomic DNA using Amplitaq Gold DNA Polymerase (Invitrogen, 4311820) and primers (F2: TCTTGTGGAAAGGACGAAACACCG; R2: TCTACTATTCTTTCCCCTGCACTGT). The resulting DNA fragment (∼ 242 bp) was gel purified. In a 2^nd^ PCR reaction illumina-compatible P7 and custom stacked barcodes (Supplement Table 2_BClist) including the standard illumina P5 forward primer were introduced into samples by PCR amplification and gel purified for the final product (∼180-200 bp). The final product was quantified by Agilent Bioanalyzer DNA High-sensitivity Assay (Agilent 5067-4626) and pooled together in equal molar ratio and analyzed by NGS. A 5% PhiX spike in used was for quality purposes. Illumina libraries were either sequenced with a 300cycle MiSeqv2 kit or a 76 cycle NextSeq 500/550 kit by single-end sequencing using NextSeq mid-output. (DRYAD doi to be generated)

### Quantification and analysis of screen data

The quantification of guides was done using a strict exact match to the forward primer, sample barcode, and guide sequence. MAGeCK was used for the identification of essential sgRNAs by running the “mageck test” command on the raw sgRNA counts. MAGeCK employs median normalization followed by a Negative Binomial modeling of the counts, and provides the log fold change (lfc) and p-values at both the guide and gene levels (Li et al., 2014).

### GFP competition and sgRNA complementation assay

TO-HCT116, TO-HCT116_mAID-ORC2^gr^, U2OS, U2OS_mAID-ORC2^gr^, HCT116 *p53^-/^*^-^, HCT116 *p53^-/-^*_mAID-ORC2^gr^, *ORC2^-/-^* p44, and *ORC2* p44^-/-^_ mAID-ORC2^gr^ cells were transduced with different sgRNA-Cas9-GFP lentivirus respectively with the MOI at 0.3-0.4 to ensure one copy of sgRNA transduction per cell. Cells were passaged every 3 days from day 3 (P1) post-transduction to day 21(P7), and at the same time, GFP percentage were evaluated by guava easyCyte™ flow cytometer. Three technical repeats were done for each datapoint. GFP percentage of day 3 (P1) for each sgRNA was normalized to 100 %, and then result of each passage was normalized to day 3 correspondingly. All experiments were repeated twice at least.

### Generating endogenous ORC2 KO mAID-ORC2^gr^ cell lines

TO-HCT116 cells were transduced with mAID-ORC2^gr^ via retroviral transduction and selected with 200 μg/ml of hygromycin to grow TO-HCT116-mAID-ORC2^gr^ cells. sgRNA_ORC2-1- epCas9-1.1-mCherry plasmid was transiently transfected into TO-HCT116-mAID-ORC2^gr^ cells using Lipofectamine 2000 Transfection Reagent (ThermoFisher #11668019) following manufacturer’s protocol. Fresh medium was replaced 6 h after transfection. Cells were harvest by 0.25% trypsin-EDTA after 24 hr., washed once with PBS, and then resuspended in sorting buffer containing 2% FBS, 2 mM EDTA, and 25mM HEPES pH7.0. Single Cell was FACS sorted by gating on mCherry fluorescence intensity into 96-well plates with 200 μl fresh medium containing 200 μg/ml of hygromycin. Single cell clone was expanded by transferring to 24-well plate, 6-well plate, and 10 cm culture dish once they reached 90% confluency.

### Cell Proliferation assays

TO-HCT116, ORC2_H-2, ORC2_H-4, and ORC2_H-5 were treated with or without 0.75µg/ml doxycycline for 24 hours before seeding. For each cell line, 150,000 cells were seeded in each well at day 1, and medium was changed every day. Every day we harvested 3 wells for each cell lines for counting. Cells were stained with 0.4 % trypan blue solution and live cells were counted by automated cell counter.

### Immunoprecipitation, Immunoblotting and quantitation

Cells were incubated with RIPA buffer (150 mM NaCl, 1 % NP-40, 0.5 % Sodium deoxycholate, 0.1 % SDS, 25 mM Tris-HCl PH 7.4) on ice for 15 minutes. Cell lysates were added with Laemmli buffer and ran on SDS-PAGE followed by western blotting to detect proteins by indicated antibodies: Primary antibodies used include anti-ORC2 (rabbit polyclonal #CS205, in-house), anti-ORC3 (rabbit polyclonal #CS1980, in-house), ani-ORC1 (mouse monoclonal #pKS1-40, in-house), anti-CDC6 (mouse monoclonal #DCS-180, EMD Millipore), anti-ATM (rabbit monoclonal #ab32420, abcam), anti-pATM(S1981) (rabbit monoclonal #ab81292, abcam), anti-CHK1 (rabbit monoclonal #ab40866, abcam), anti-pCHK1(S345) (rabbit polyclonal #2348, Cell Signaling), anti-pCHK2(T68) (rabbit polyclonal #2197, Cell Signaling), anti-ATR (rabbit polyclonal #ab2905, abcam), anti-pATR(T1989) (rabbit polyclonal #ab227851, abcam), anti-pATR(S428) (rabbit polyclonal #2853, Cell Signaling), anti-p-gH2AX(S139) (rabbit monoclonal #9718, Cell Signaling), anti-b-Actin (mouse monoclonal #3700, Cell Signaling). Secondary antibodies include ECL^TM^ anti-Rabbit IgG Horseradish Peroxidase linked whole antibody (#NA934V, GE Healthcare) and ECL^TM^ anti-mouse IgG Horseradish Peroxidase linked whole antibody (#NA931V, GE Healthcare).

Relative ORC2 (or mAID-ORC2^gr^) and ORC3 Protein level in each cell line was quantitated by normalizing band intensity to β-Actin of each cell line and then eventually normalized to HCT116 cells using ImageJ software.

### Cell cycle analysis and pulse EdU label

In double-thymidine block and release experiment, cells were first incubated with 2 mM thymidine for 18 hours. After 3 times PBS wash, cells were released into fresh media with or without (0.75 µg/ml) doxycycline for 9 hours. Next, 2 mM thymidine were added into the media for 16 hours. 500 nM of auxin was added into the medium if needed 4.5 hr. before released. When released from thymidine block, 0 hr. time point cells were harvest, and the rest were washed with PBS for 3 times and released into fresh medium ±dox and auxin. Cells were collected at indicated time point and 10 µM EdU were added into the medium 2 hours before each harvest (Including time 0). Cells were fixed and processed following Click-iT™ EdU Alexa Fluor™ 488 Flow Cytometry Assay Kit manufacturer’s manual (ThermoFisher #C10420) and DNA was stained with FxCycle^TM^ Violet Stain (ThermoFisher #F10347).

### Mitotic index flow cytometry

TO-HCT116 and ORC2_H-2 cells were pre-treated with doxycycline for 24 hr. when needed in this experiment. Cells were trypsinized and harvest at different time points after auxin treatment, and immediately fixed with 4 % Paraformaldehyde in PBS for 15 min, centrifuged at 1000 xg for 7 min. to remove fixation and washed with 1 % BSA-PBS and centrifuged. Next, cells were permeabilized with 0.5 % triton x-100 in 1 % BSA-PBS for 15 min. at room temperature, centrifuged, and washed with 1% BSA-PBS and centrifuged. Next, primary antibody anti- pH3S10 antibody (mouse monoclonal #9706, Cell Signaling) were incubated at 37 °C for 45 min. Cells were then washed 3 times in 1 % BSA-PBS +0.1 % NP-40, and incubated with secondary antibody (Donkey anti-Mouse Alexa Fluor 647 #715-605-151 Jackson ImmunoResearch) at 37 °C for 50 min. in the dark. Lastly, after 3 washes cells were incubated with FxCycle^TM^ Violet Stain (ThermoFisher). The positive/negative gates for pH3S10 were gated on a negative control, which is unstained cells.

### Cell extraction and MCM2 flow cytometry

EdU-pulse-labeled asynchronous TO-HCT116, ORC2_H-2, ORC2_H-5 cells with or without doxycycline and auxin treatment were harvest, washed with PBS, and processed based on the protocol from (Matson et al., 2017) with slight optimization. In brief, for non-extracted cells, cells were fixed with 4 % Paraformaldehyde in PBS for 15 min, and then centrifuged at 1000 xg for 7 min. to remove fixation and washed with 1 % BSA-PBS and centrifuged. Next, cells were permeabilized with 0.5 % triton x-100 in 1 % BSA-PBS for 15 min, centrifuged, and washed with 1% BSA-PBS and centrifuged. For chromatin extracted cells, cells were lysed on ice for 5 min. in CSK buffer (10mM PIPES/KOH pH 6.8, 100 mM NaCl, 300 mM sucrose, 1 mM EGTA, 1 mM MgCl2, 1 mM DTT) with 0.5 % triton x-100 with protease and phosphatase inhibitors. Cells were centrifuged and washed with 1 % BSA-PBS twice and then fixed in 4% paraformaldehyde in PBS for 15 min. After one wash with PBS, cells were processed following Click-iT™ EdU Alexa Fluor™ 488 Flow Cytometry Assay Kit manufacturer’s manual (ThermoFisher #C10420), but instead of saponin-based permeabilization and wash reagent, we used 1 % BSA-PBS +0.1 % NP-40 for all washing steps. Next, cells were incubated with Anti-MCM2 antibody (mouse monoclonal #610700, BD Biosciences) at 37 °C for 40 min. in the dark. Cells were then washed 3 times in 1 % BSA-PBS +0.1 % NP-40, and incubated with secondary antibody (Donkey anti-Mouse Alexa Fluor 647 #715-605-151 Jackson ImmunoResearch) at 37 °C for 50 min. in the dark. Finally, cells were washed for 3 times and incubated with FxCycle^TM^ Violet Stain (ThermoFisher). The positive/negative gates for MCM were gated on a negative control, which is unstained cells.

### Immunofluorescence Staining

TO-HCT116, ORC2_H-2, ORC2_H-4, and ORC2_H-5 cells were grown on coverslips for 2 days with or without doxycycline and auxin treatment. When harvest, coverslips were transferred to 6- well plate and rinse with PBS. Cells were fixed in 4 % Paraformaldehyde for 10 min. at room temperature. Next, cells were washed three times for 5 min. in cold PBS. Cells were then permeabilized in 0.5 % Triton x-100 -PBS for 9 min. After three PBS wash, cells were blocked with 5 % normal goat serum (NGS) -PBS +0.1 % Tween for 1 hr. For primary antibody incubation, antibodies were diluted in 1 % NGS-PBS +0.1 % Tween and incubated for 17 hr. in the cold room. Primary antibodies used include anti-CENP-C antibody (Mouse monoclonal #ab50974, Abcam), anti-pCHK1(S345) (rabbit polyclonal #2348, Cell Signaling), anti-p- gH2AX(S139) (rabbit monoclonal #9718, Cell Signaling), and anti-pATM(S1981) (Mouse monoclonal #ab36180, Abcam). Cells were washed three times for 5 min. in 1 % NGS-PBS +0.1 % Tween before 1 hr. secondary antibody incubation at room temperature. Secondary antibodies used are Goat Anti-Mouse IgG H&L Alexa Fluor® 647 (#ab150115, Abcam) and Goat Anti-Rabbit IgG H&L Alexa Fluor® 488 (#ab150077, Abcam). Next, cells were stained with 1 µg/ml DAPI and mounted with VECTASHIELD® Antifade Mounting Medium (#H-1000-10, Vector Laboratories). Images were taken using a Perkin Elmer spinning disc confocal equipped with a Nikon-TiE inverted microscope using 60X objective oil lens with an Orca ER CCD camera. Images presented are maximum intensity projections of a z-stack (z=0.3µM).

To study the nuclear and cellular morphology HCT116 p53 WT and p53 null cells and *ORC1^-/-^ (*B14) and *ORC2^-/-^* (P44) cells were plated on coverslips. On day 2 cell were fixed with 4% PFA and the method described above was followed. Primary antibody against Lamin B1 (Abcam ab16048) was used as a marker for nuclear envelope. Secondary antibody used is Goat Anti-Rabbit IgG Alexa Fluor® 594 (Abcam ab150084). In addition, Phalloidin iFluor® 488 (Abcam ab176753) was used to stain for cytoskeleton and DNA was detected with 1µg/ml Hoechst dye (ThermoFisher #62249). Mounted coverslips were imaged with Perkin Elmer spinning disc confocal equipped with a Nikon-TiE inverted microscope using 40X objective lens with an Orca ER CCD camera. Images presented are single channel average intensity projections or merged multi-channel maximum intensity projections of z-stacks.

### ORC2 Nuclear volume quantitation

Cells nuclear were fixed and stained with DR™ Fluorescent Probe Solution as per the manufacturer’s guidelines (ThermoFisher #62251). Images were taken using a Perkin Elmer spinning disc confocal equipped with a Nikon-TiE inverted microscope using 60X objective lens with an Orca ER CCD camera. Images presented are maximum intensity projections of a z-stack (z=0.3µM). Nuclear size was analyzed with volocity software (version 6.3.1).

### Live cell microscopy

TO-HCT116 and ORC2_H-2 cells were seeded in ibidi µ-Slide 8 Well Glass Bottom and in the presence or absence of 0.75 µg/ml doxycycline for 24 hours. Next, 2 mM thymidine were added and incubated ±dox for 24 hours. Two hours prior to washing out thymidine, 500 nM auxin were added to the dox treated wells. Samples were then imaged starting at 4 hours after thymidine release and the timepoints reconstructed as from a movie using volocity software. Images were taken using a Perkin Elmer spinning disc confocal equipped with a Nikon-TiE inverted microscope using 40X objective lens with an Orca ER CCD camera. Images presented are maximum intensity projections of a z-stack (z=3 µM). Frames were taken approximately every 5 minutes.

### Quantitative PCR

Total RNA of HCT116 *p53^-/-^* and *ORC2^-/-^* cells were extracted using Rneasy Mini Kit (Qiagen #74104) following manufacturer’s handbook and quantified by Nanodrop (ThermoFisher). RNA was then converted to cDNA by doing reverse transcription using oligo(dT) or random hexamer primers provided by TaqMan Reverse Transcription Reagents (#N8080234, Applied Biosystems). Primer pairs for quantitative PCR are designed to PCR exon-exon junction (Supplement Table 3_exon primers) and each PCR was done in triplicates. The delta-Ct (ΔCt) value was obtained from subtracting Actin mean Ct from each primer pair mean Ct in each cell line. The delta-delta-Ct (ΔΔCt) value was calculated by subtracting HCT116 *p53^-/-^* ΔCt from ORC2^-/-^ ΔCt for each primer pair individually. Fold change (FC) for each primer pair in *ORC2^-/-^* cells compared to HCT116 p53^-/-^ cells was calculated as FC = 2 (to the power of ΔΔCt).

### Transmission electron microscopy

HCT116 and *ORC1^-/-^* cells were harvest and washed twice with PBS. Cells were pelleted and resuspended in 1 mL of 2.5 % glutaraldehyde in 0.1 M sodium cacodylate solution (pH 7.4) overnight at 4 °C. Fixative was removed, and in each step, 200 µl of the solution was left in the tubes. Pellet was washed with 0.1 M sodium cacodylate buffer. Next, 4 % Melt agarose solution was added to the tube and centrifuged immediately at 1,000 x g for 10 min. at 30 °C, and then directly transferred the tube to 4 °C or ice for 20 min. to solidify the agarose. Agarose was washed twice with 0.1 M cacodylate buffer. Next, 1 % osmium tetraoxide (OsO4) solution was added and let stand for 1 hr. followed by three 0.1 M cacodylate buffer washes. Samples were then dehydrated by a graded ethanol series (50 %, 60 %, 70 %, 80 %, 90 %, 95 %, 100 %, respectively). Finally, samples were embedded in 812 Embed resin and sectioned in 60-90 µM using Ultramicrotome. Hitachi H-7000 Transmission Electron Microscopy was used to image the sample.

### Nanopore Long read sequencing and analysis

High molecular weight DNA was isolated using the MagAttract kit (Qiagen # 67563). The quality of the DNA from the was assessed on femtopulse (Agilent) to ensure DNA fragments were >40kb on average DNA was sheared to 50kb via Megarupter (diagenode). After shearing, the DNA was size selected with a SRE kit (Circulomics) to reduce the fragments <20kb. After size selection, the DNA under when a-tailing and damage repair followed by ligation to sequencing specific adapters. The ½ prepared library was mixed with library loading beads and loaded on to a PROM-0002 flow-cell and was allowed to sequence for 24 hours. After 24 hours the flow- cell was treated with DNase to remove stalled DNA followed by a buffer flush. The second ½ of the library was then loaded and allowed to sequencing for 36 hours. The DNA was base called via Guppy 3.2 in High accuracy mode. Long reads were aligned to the reference human genome using NGMLR (https://github.com/philres/ngmlr) and structural variants were identified using Sniffles (https://github.com/fritzsedlazeck/Sniffles) (Sedlazeck et al., 2018). The alignments and structural variants were then visualized using IGV (https://igv.org/).

## Acknowledgements

We thank Dr. Leemor Joshua-Tor for comments on the manuscript and Dr. Anindya Dutta for providing cell lines. We also thank Jennifer Shapp for technical assistance. This work was supported by grants from the National Cancer Institute (P01- CA13106 and a Cancer Center Support Grant P50-CA045508.

## Author Contributions

All the experiments were performed by H-C.C. and K.B., microscopy was performed in collaboration with H.A., the computational analysis of genomic data was done in collaboration with O.E.D., the CRISPR-Cas9 genome tiling methods and library preparation were done in collaboration with O.K., K.H. K.C. and C.V. and the genomic DNA sequencing was done in collaboration with R.W.M. and data analyzed by S.A. and M.S. Genomic copy number determination was done with P.A.. Experiments were designed and analyzed by H-C.C, K.B, O.K, C.V. and B.S. H-C.C, K.B. and B.S. wrote the paper.

---

**Figure 1—Figure supplement 1.**
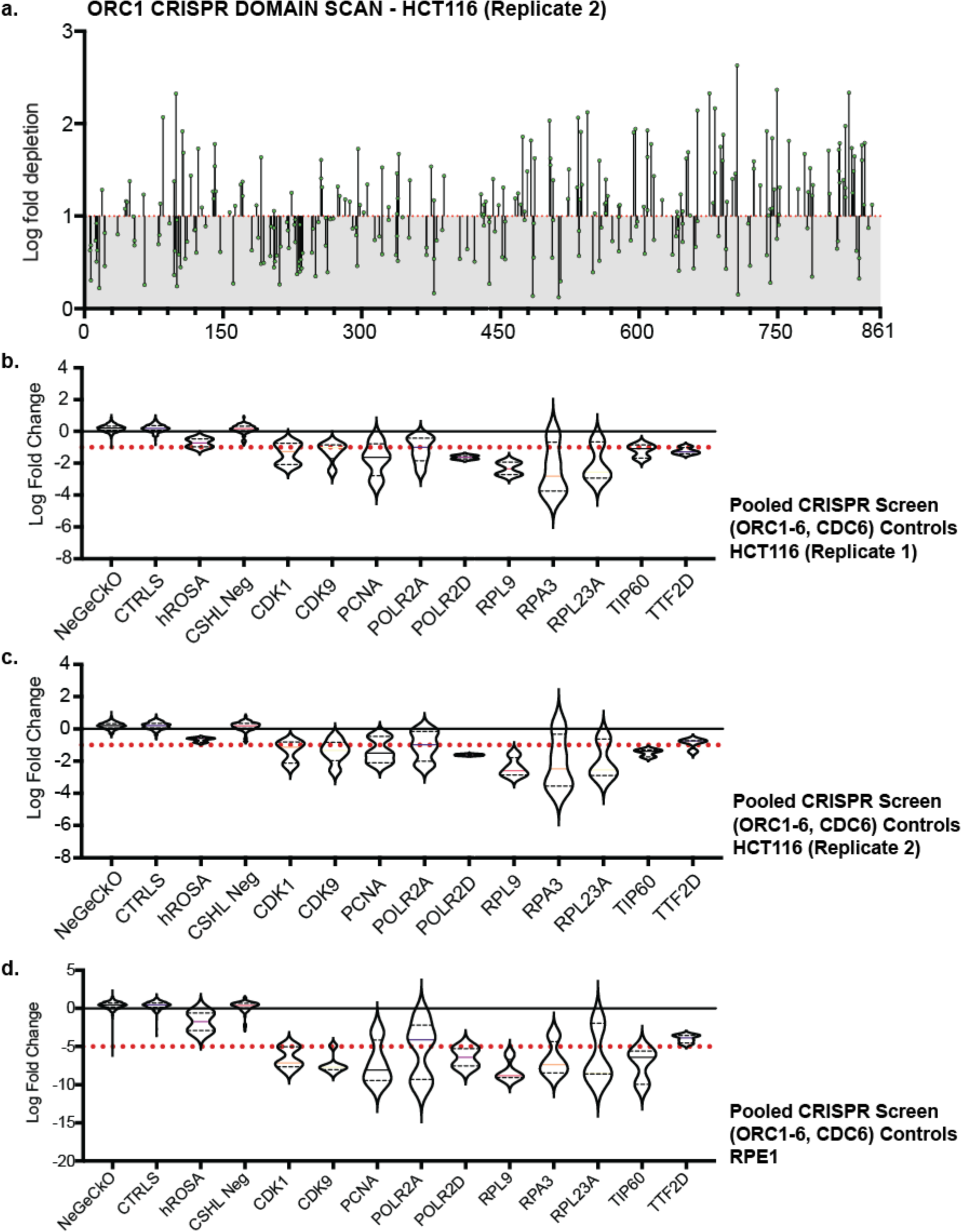
Tiling sgRNA CRISPR screen data and controls. (a) tiling sgRNA map of ORC1 (replicate 2) in HCT116. Mapped as Log fold depletion (inverted LFC scale) as calculated by MaGeCK (Li et al., 2014) on y axis vs the codon that is disrupted by the guide RNA on the x axis. Effect of guide RNA is interpreted as essential if its depletion is more than 1 log fold (red dotted line). (b) tiling sgRNA controls for HCT116 (replicate 1). Violin plots mapped as distribution of Log fold depletion (MaGeCK) for each guide RNA from for negative (NeGeCKO, CTRLS, hROSA, and CSHL-neg library) or positive control (CDK1, CDK9, PCNA, POLR2A, POLR2D, RPL9, RPA3, RPL23A, TIP60, TTF2D) subset. The median and quartiles of LFC for each subset are indicated within each violin plot. Cut-off of essentiality is LFC ≥ -1, indicated by red dotted line (Highest log fold depletion value of all negative controls and more than the median of positive controls. (c) tiling sgRNA controls for HCT116 (replicate 2). (d) tiling sgRNA controls for RPE-1. Cut-off of essentiality is LFC ≥ -5, indicated by red dotted line.

**Figure 1—Figure supplement 2.**
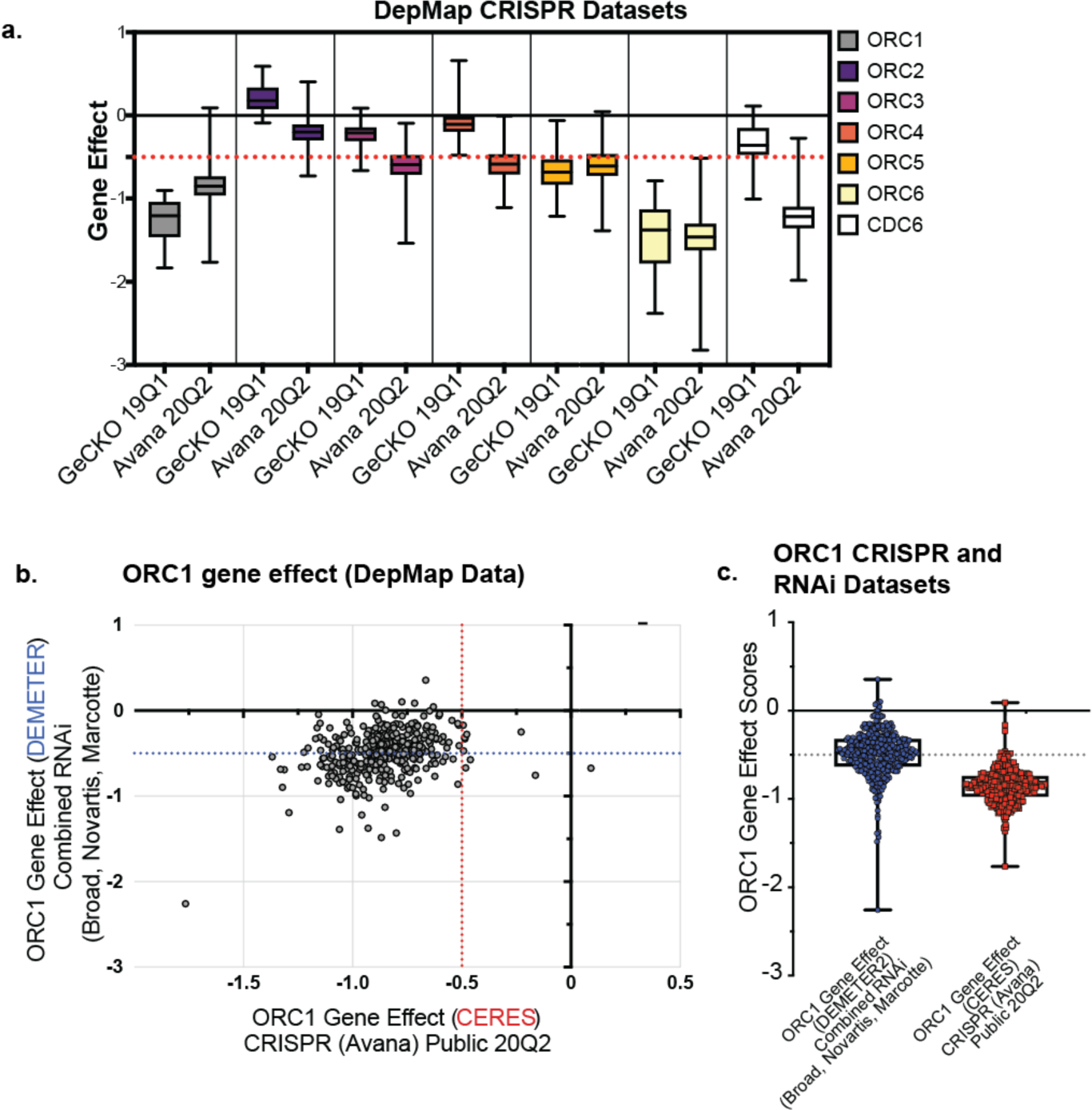
DepMap analyses of *ORC1* data. (a) Distribution of Gene Effect scores of *ORC1-6* and *CDC6* across all the cell lines used in either the GeCKO 19Q1 or Avana 20Q2 CRISPR screens reported on DepMap [https://doi.org/10.6084/m9.figshare.7668407, https://doi.org/10.6084/m9.figshare.12280541.v4; (Meyers et al., 2017b)]. Each box plot represents gene effect range displayed in the tested cell lines. The red dotted line represents the gene effect score below which genes are scores as essential. (b) *ORC1* gene effect values for CRISPR (CERES; (Meyers et al., 2017b) vs RNAi (DepMap, 2019; McFarland et al., 2018)mapped as xy scatter for ∼390 common cell lines used in the screens. Red dotted line bifurcates the plot at CRISPR based gene effect score of less than -0.5 is considered essential to cell line. Blue dotted line bifurcates the plot at RNAi based gene effect score of less than -0.5 is considered essential to cell line. (c) Distribution of *ORC1* gene effect scores across all the cell lines used in CRISPR Avana 20Q2 and RNAi datasets respectively ((McFarland et al., 2018; Meyers et al., 2017b).

**Figure 2—Figure supplement 1.**
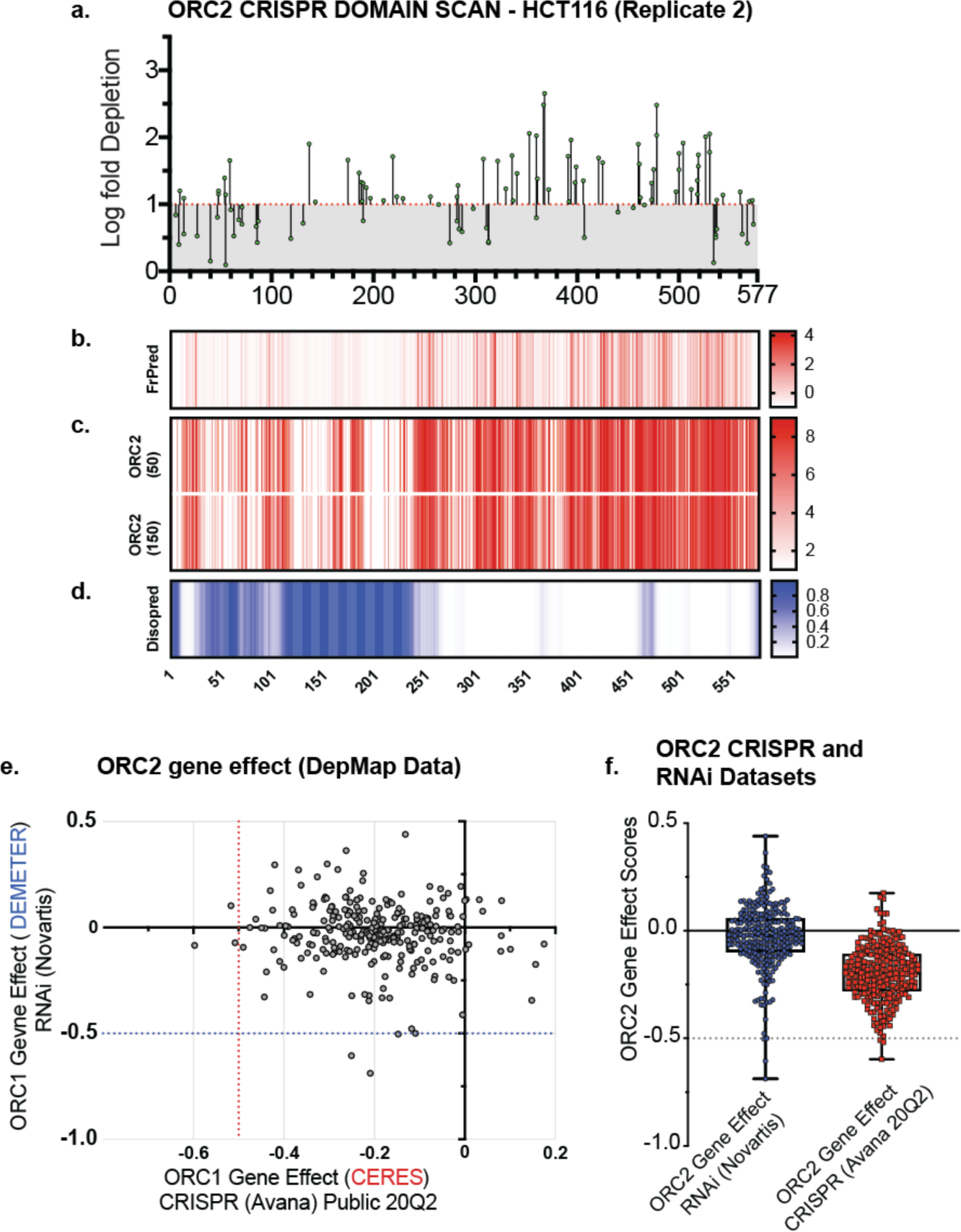
(a) Tiling sgRNA map of ORC2 (replicate 2) in HCT116. (b) FrPred (https://toolkit.tuebingen.mpg.de/frpred) of hORC2 (NP_006181.1) shown as gradient heat map of conservation score vs amino acid position. (d) Consurf (https://consurf.tau.ac.il/) of hORC2 – (upper) ORC2 (50) subset (50 HMMER Homologues collected from UNIREF90 database, Max sequence identity = 95%, Min sequence identity 50, Other parameters = default), and (lower) ORC2 (150) subset (150 HMMER Homologues collected from UNIREF90 database, Max sequence identity = 95%, Min sequence identity 35, Other parameters = default). Data represented as heatmap of Conservation scores of each amino acid position. (e) Disopred (http://bioinf.cs.ucl.ac.uk/psipred/) plot of hORC2 – heatmap representing amino acids within intrinsically disordered regions of the protein.

**Figure 2—Figure supplement 2.**
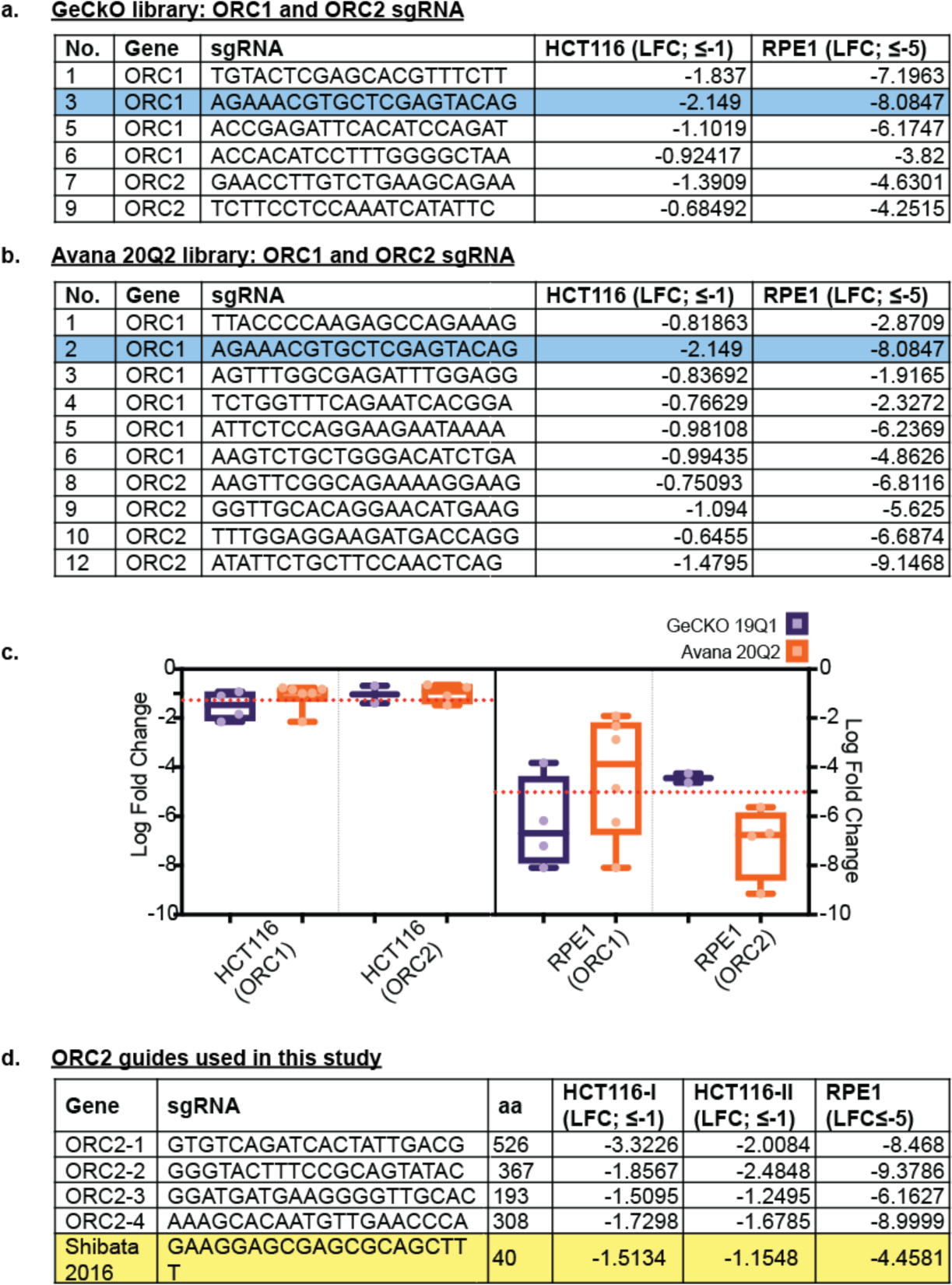
(a) Table listing guide RNA sequences used in GeCKO 19Q1 library against *ORC1* and *ORC2* that were also present in our tiling sgRNA screen; columns HCT116(LFC ≤ -1) and RPE1(LFC ≤-5) show the LFC of the GeCKO guides in our screen. (b) Table listing guide RNA sequences used in Avana 20Q2 library against *ORC1* and *ORC2* that were also present in our tiling sgRNA screen; columns HCT116(LFC ≤ -1) and RPE1(LFC ≤-5) show the LFC of the GeCKO guides in our screen. Rows highlighted blue represent the only common guide against *ORC1* used between GeCKO and Avana screens (c) Distribution of the LFC values for the GeCKO and Avana guides in HCT116 and RPE-1 (graphical representation of the tables (a) & (b). (d) *ORC2* guides selected for single guide studies. Row highlighted yellow – guide RNA sequence used in the Shibata et. Al 2016 study.

**Figure 2—Figure supplement 3.**
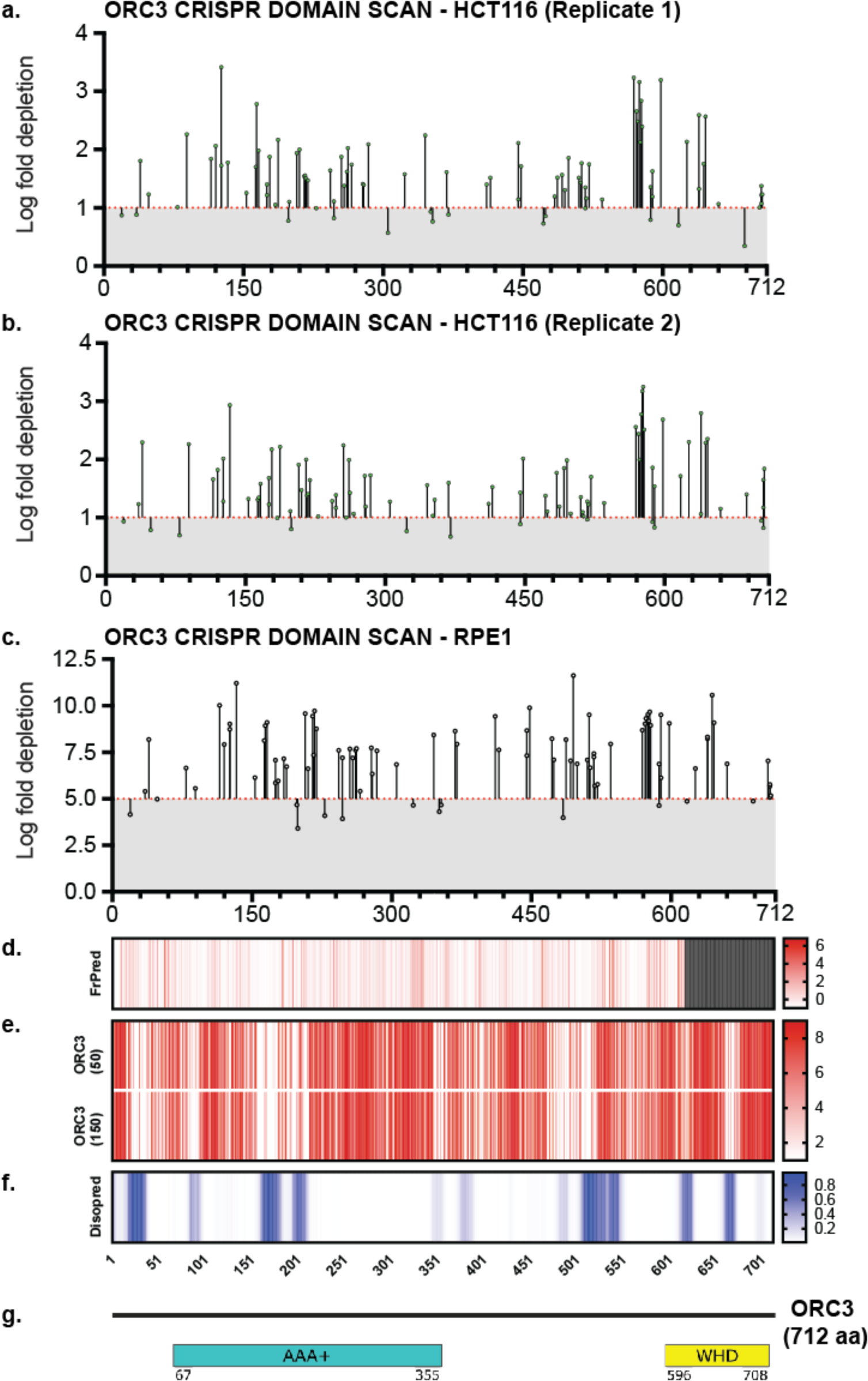
Tiling sgRNA CRISPR screen data contd. (a) tiling sgRNA map of ORC3 (replicate 1) in HCT116. Mapped as Log fold depletion as calculated by MaGeCK on y axis vs the codon that is disrupted by the guide RNA on the x axis. Effect of guide RNA is interpreted as essential if its depletion is more than 1 log fold (red dotted line). (b) tiling sgRNA map of ORC3 for HCT116 (replicate 2). (c) tiling sgRNA map of ORC3 for RPE-1 cell line. Cutt- off of essentiality is LFC ≥ -5, indicated by red dotted line. (d) FrPred (https://toolkit.tuebingen.mpg.de/frpred) of hORC3 (NP_862820.1) shown as gradient heat map of conservation score vs amino acid position. (e) Consurf (https://consurf.tau.ac.il/) of hORC3 – (upper) ORC3 (50) subset (50 HMMER Homologues collected from UNIREF90 database, Max sequence identity = 95%, Min sequence identity 50, Other parameters = default), and (lower) ORC3 (150) subset (150 HMMER Homologues collected from UNIREF90 database, Max sequence identity = 95%, Min sequence identity 35, Other parameters = default). Data represented as heatmap of Conservation scores of each amino acid position. (f) Disopred (http://bioinf.cs.ucl.ac.uk/psipred/) plot of hORC3 – heatmap representing amino acids within intrinsically disordered regions of the protein. (g) Schematic of domain architecture of ORC3.

**Figure 2—Figure supplement 4.**
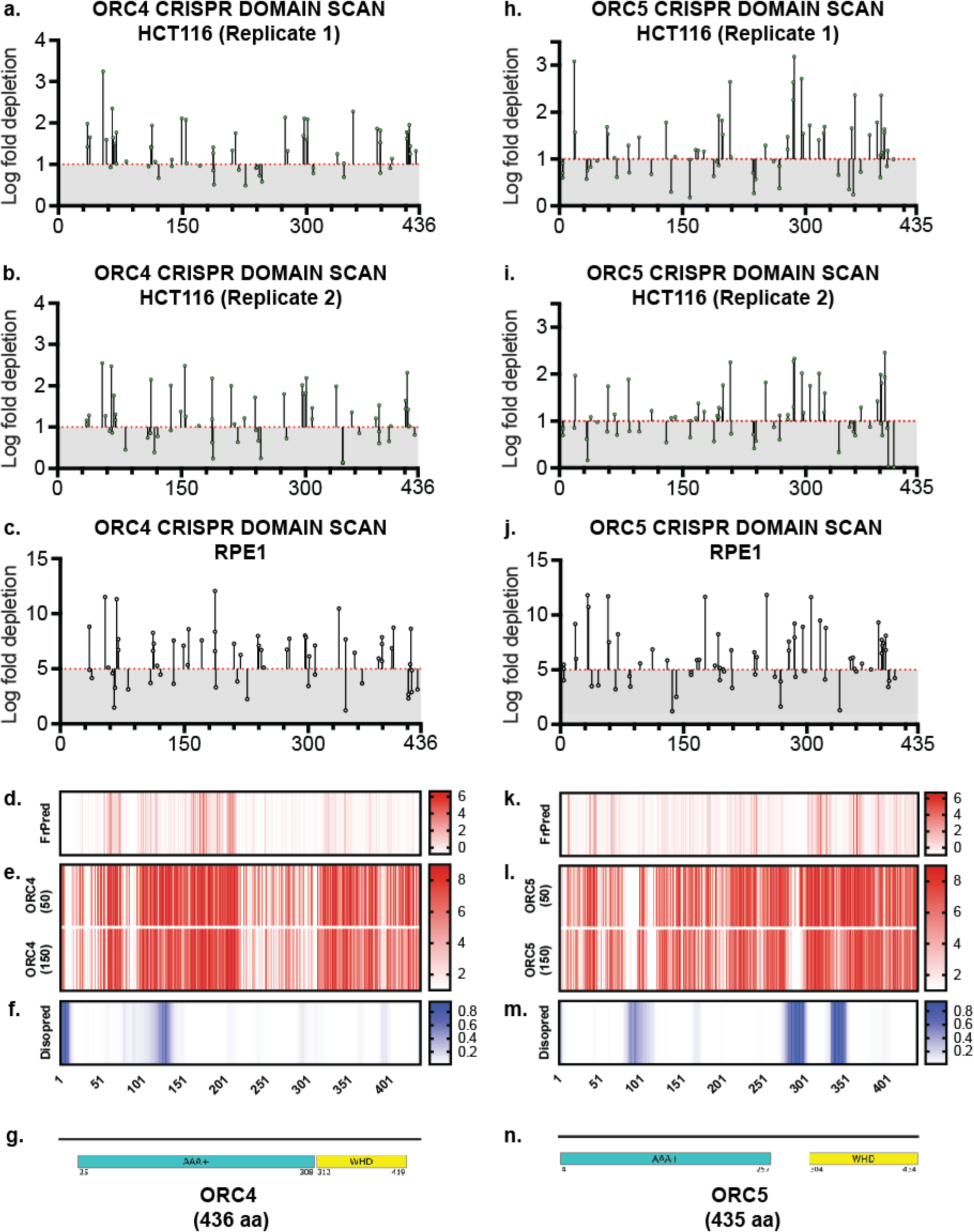
Tiling sgRNA CRISPR screen data contd. (a) tiling sgRNA map of ORC4 (replicate 1) in HCT116. Mapped as Log fold depletion as calculated by MaGeCK on y axis vs the codon that is disrupted by the guide RNA on the x axis. Effect of guide RNA is interpreted as essential if its depletion is more than 1 log fold (red dotted line). (b) tiling sgRNA map of ORC4 for HCT116 (replicate 2). (c) tiling sgRNA map of ORC4 for RPE-1 cell line. Cutt- off of essentiality is LFC ≥ -5, indicated by red dotted line. (d) FrPred (https://toolkit.tuebingen.mpg.de/frpred) of hORC4 (NP_859525.1) shown as gradient heat map of conservation score vs amino acid position. (e) Consurf (https://consurf.tau.ac.il/) of hORC4 – (upper) ORC4 (50) subset (50 HMMER Homologues collected from UNIREF90 database, Max sequence identity = 95%, Min sequence identity 50, Other parameters = default), and (lower) ORC4 (150) subset (150 HMMER Homologues collected from UNIREF90 database, Max sequence identity = 95%, Min sequence identity 35, Other parameters = default). Data represented as heatmap of Conservation scores of each amino acid position. (f) Disopred (http://bioinf.cs.ucl.ac.uk/psipred/) plot of hORC4 – heatmap representing amino acids within intrinsically disordered regions of the protein. (g) Schematic of domain architecture of ORC4. (h) tiling sgRNA map of ORC5 (replicate 1) in HCT116. (i) tiling sgRNA map of ORC5 for HCT116 (replicate 2). (j) tiling sgRNA map of ORC5 for RPE-1 cell line. (k) FrPred (https://toolkit.tuebingen.mpg.de/frpred) of hORC5 (NP_002544.1) shown as gradient heat map of conservation score vs amino acid position. (e) Consurf (https://consurf.tau.ac.il/) of hORC5 – (upper) ORC5 (50) subset and (lower) ORC5 (150) subset. Data represented as heatmap of Conservation scores of each amino acid position. (f) Disopred (http://bioinf.cs.ucl.ac.uk/psipred/) plot of hORC5 – heatmap representing amino acids within intrinsically disordered regions of the protein. (g) Schematic of domain architecture of ORC5.

**Figure 2—Figure supplement 5.**
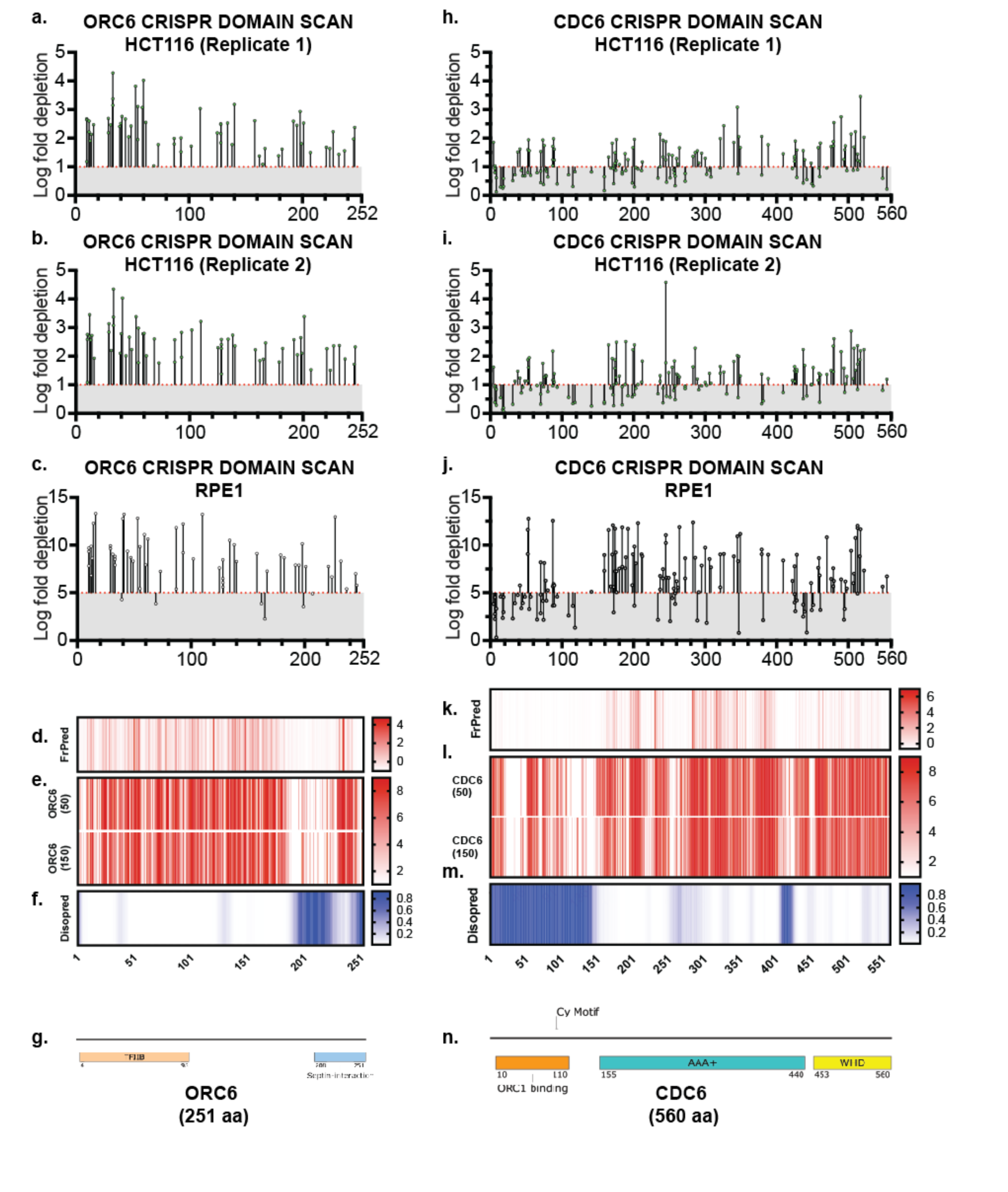
Tiling sgRNA CRISPR screen data contd. (a) Tiling sgRNA map of ORC6 (replicate 1) in HCT116. Mapped as Log fold depletion as calculated by MaGeCK on y axis vs the codon that is disrupted by the guide RNA on the x axis. Effect of guide RNA is interpreted as essential if its depletion is more than 1 log fold (red dotted line). (b) Tiling sgRNA map of ORC6 for HCT116 (replicate 2). (c) Tiling sgRNA map of ORC6 for RPE-1 cell line. Cutt- off of essentiality is LFC ≥ -5, indicated by red dotted line. (d) FrPred (https://toolkit.tuebingen.mpg.de/frpred) of hORC6 (NP_055136.1) shown as gradient heat map of conservation score vs amino acid position. (e) Consurf (https://consurf.tau.ac.il/) of hORC6 – (upper) ORC6 (50) subset and (lower) ORC6 (150) (f) Disopred (http://bioinf.cs.ucl.ac.uk/psipred/) plot of hORC6-heatmap representing amino acids within intrinsically disordered regions of the protein. (g) Schematic of domain architecture of ORC6. (h) tiling sgRNA map of CDC6 (replicate 1) in HCT116. (i) Tiling sgRNA map of CDC6 for HCT116 (replicate 2). (j) Tiling sgRNA map of CDC6 for RPE-1 cell line. (k) FrPred (https://toolkit.tuebingen.mpg.de/frpred) of hCDC6 (NP_001245.1) shown as gradient heat map of conservation score vs amino acid position. (e) Consurf (https://consurf.tau.ac.il/) of hCDC6 – (upper) CDC6 (50) subset and (lower) ORC5 (150) subset. Data represented as heatmap of Conservation scores of each amino acid position. (f) Disopred (http://bioinf.cs.ucl.ac.uk/psipred/) plot of hCDC6 – heatmap representing amino acids within intrinsically disordered regions of the protein. (g) Schematic of domain architecture of hCDC6.

**Figure 3—Figure supplement 1.**
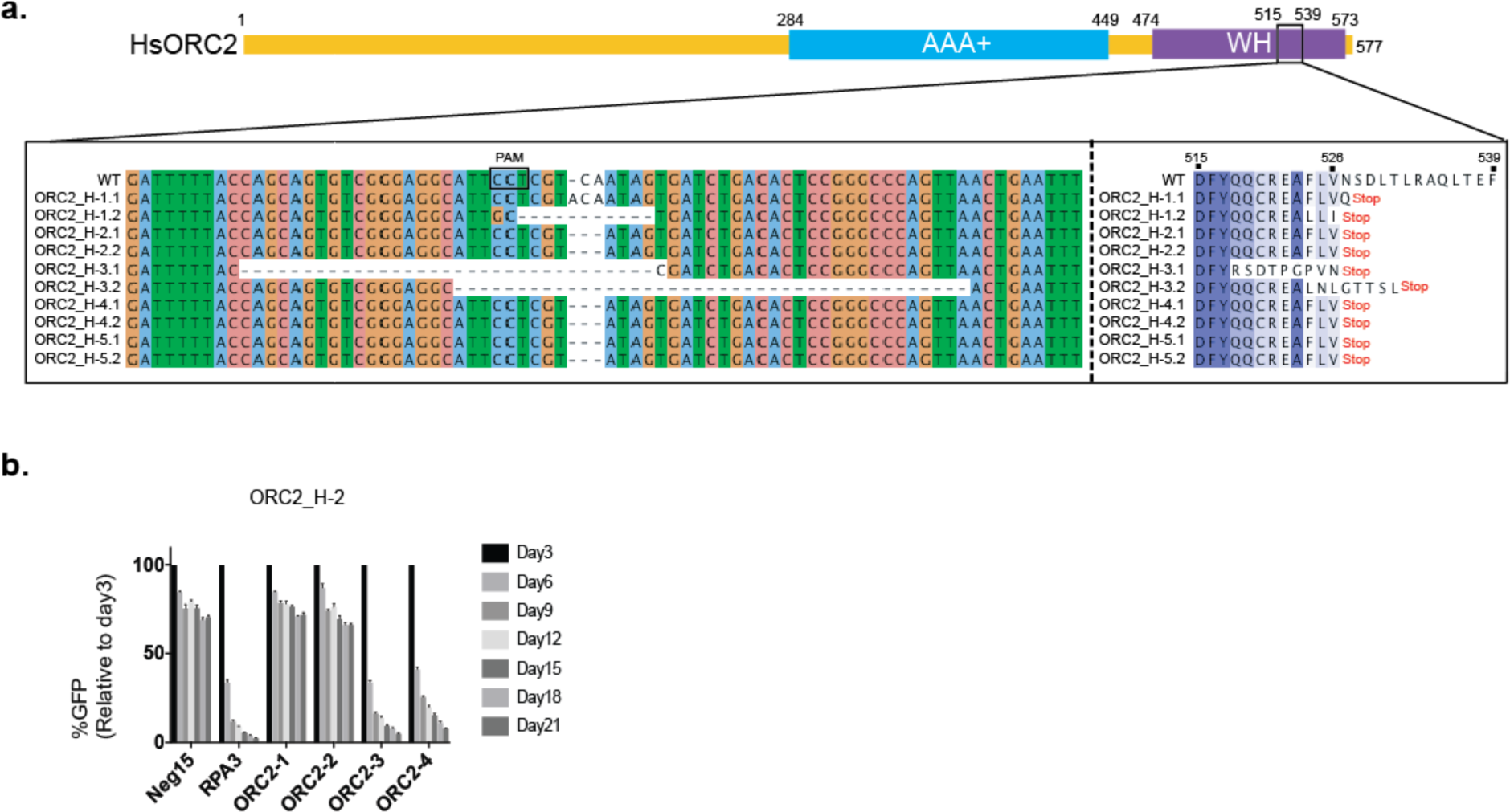
CRISPR/Cas9 ORC2-1 sgRNA mutagenesis in five cell lines. (a) Nucleotide and amino acid alignments near the sgRNA targeting site in parental TO-HCT116 and five cloned ORC2 KO cell lines. (b) ORC2_H-2 is resistant to ORC2_1 and ORC2_2 sgRNAs. Negative-selection time course assay that plots the percentage of GFP^+^ cells over time following transduction with the indicated sgRNAs with Cas9. The GFP positive percentage was normalized to the Day3 measurement. N = 3. Error bars, mean ± SD.

**Figure 3—Figure supplement 2.**
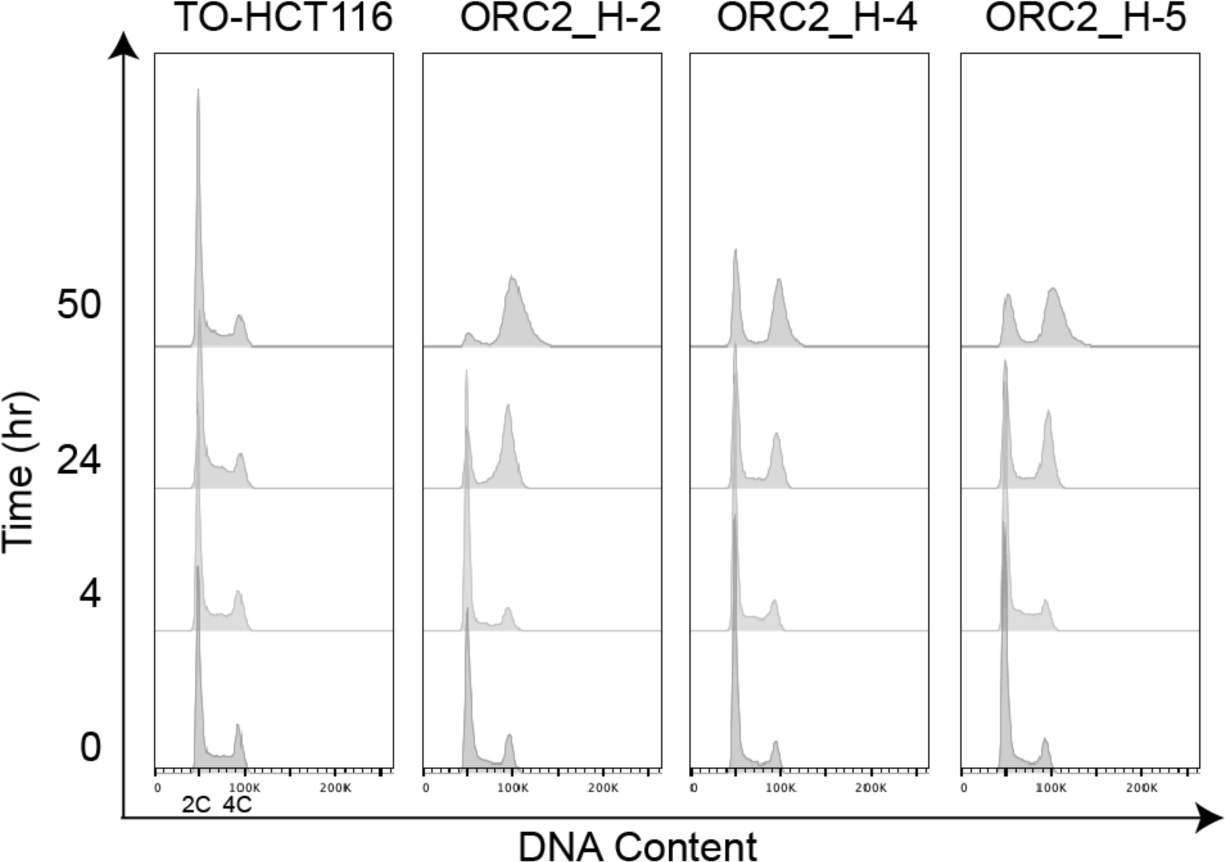
Flow cytometry analysis of cell cycle after dox and auxin treatment in TO-HCT116, ORC2_H-2, ORC2_H-4, and ORC2_H-5 cell lines. X axis refers to DNA content. Y axis represents the period of time (hr) for auxin treatment.

**Figure 3—Figure supplement 3.**
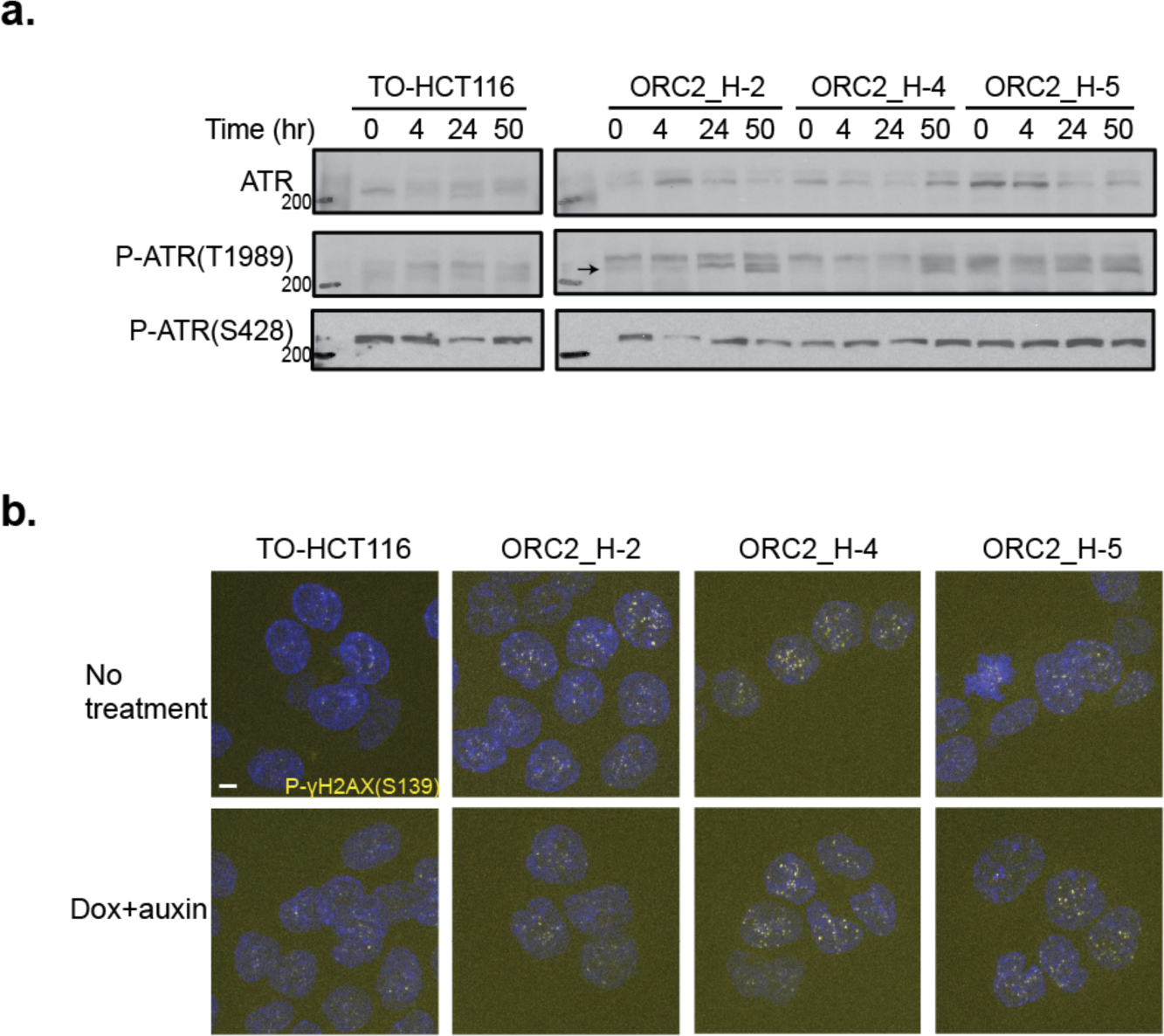
DNA damage checkpoint was activated in ORC2_H-2, ORC2_H-4, and ORC2_H-5 cell lines. (a) ATR(T1989) was phosphorylated in ORC2_H-2, -4, -5 cell lines after dox and auxin treatment for 50 hours. There was no change in p-ATR(S428) level. (b) Phosphorylation of gH2AX(S139) was seen in ORC2_H-2, -4, -5 cell lines either in the absence or presence of dox and auxin.

**Figure 5—Figure supplement 1.**
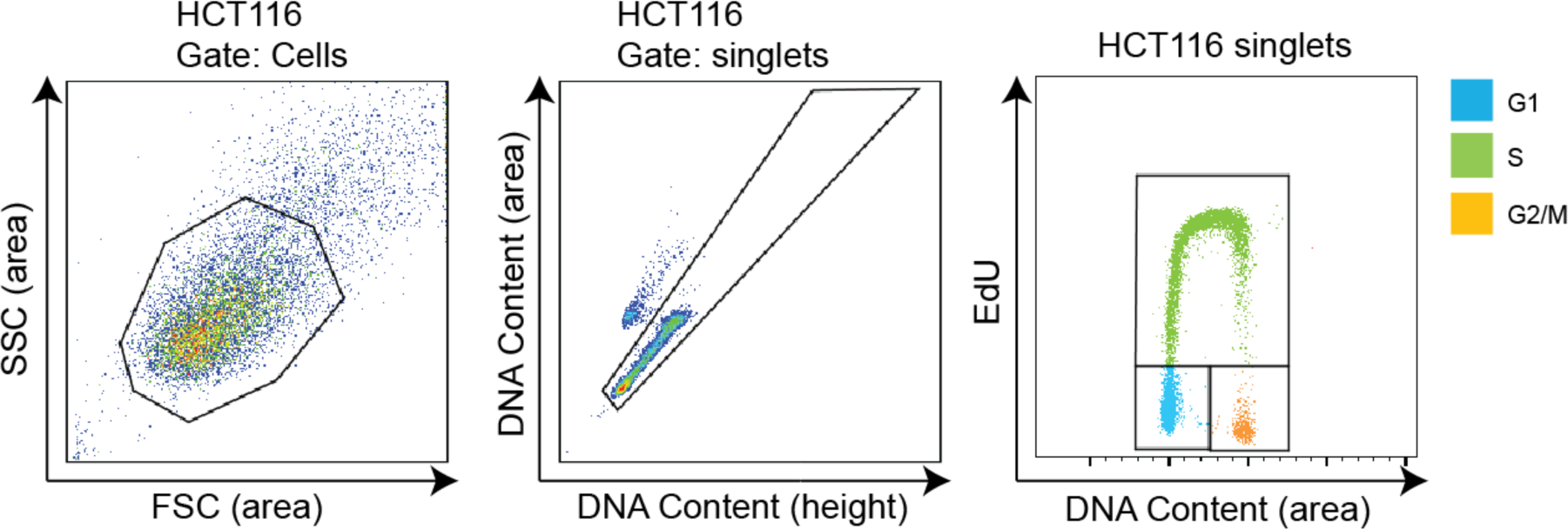
Flow cytometry gating. Example of flow cytometry gating for Figure 5b and c. First of all, FSC (area) vs SSC (area) gating was used to exclude the cell debris. Next, singlets were gated on FxCycle™ Violet (DNA content) height vs area. Cell population in G1, S, and G2/M phase was gated on the FxCycle™ Violet (DNA content) area vs EdU plot.

**Figure 6—Figure supplement 1.**
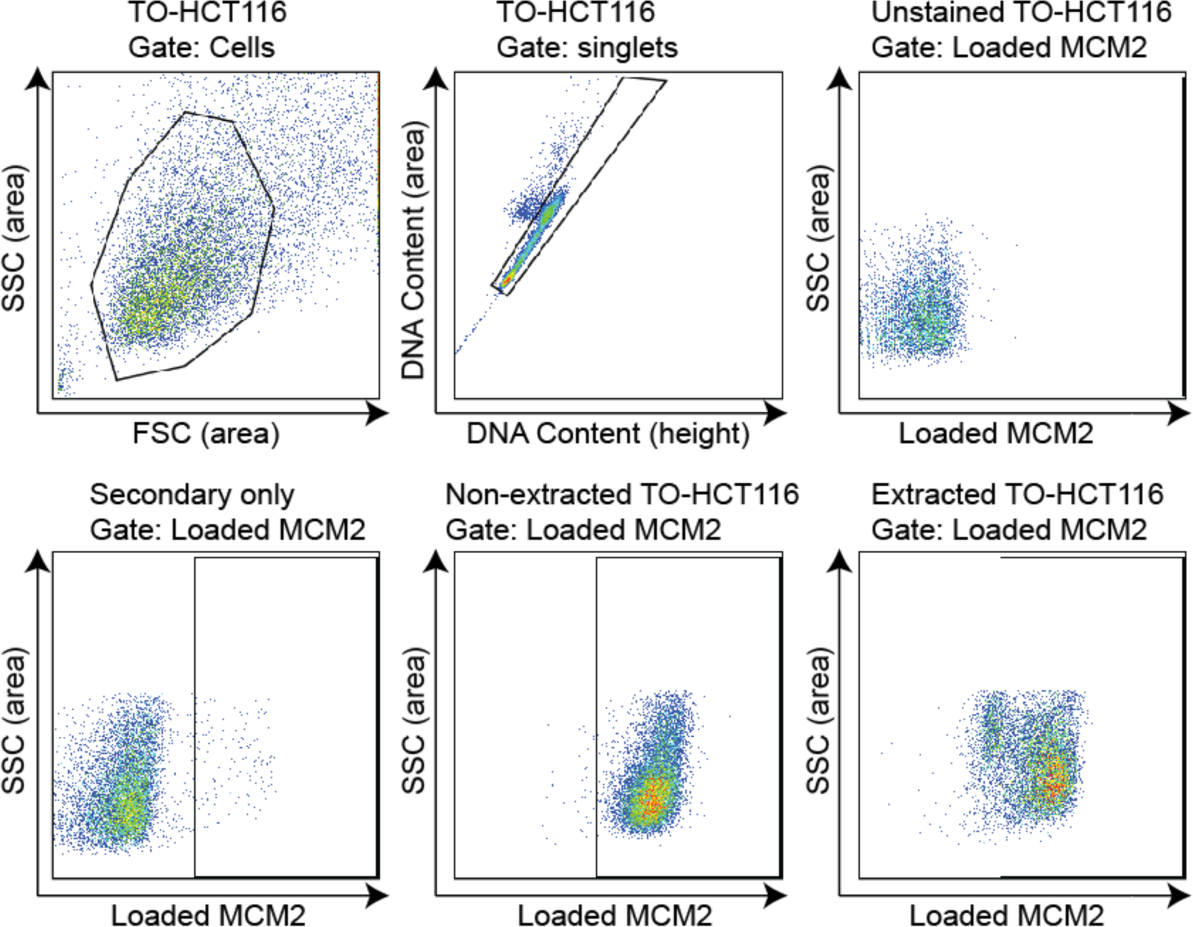
Flow cytometry gating. Example of flow cytometry gating for Figure 6. FSC (area) vs SSC (area) gating was used to exclude the cell debris. Next, singlets were gated on FxCycle™ Violet (DNA content) height vs area. MCM2 positive population was gated on the loaded MCM2 (area) vs SSC (area) of the unstained negative control. Cells stained for secondary Donkey anti-Mouse Alexa Fluor 647 antibody only showed minimum background for loaded MCM2.

**Figure 7—Figure supplement 1.**
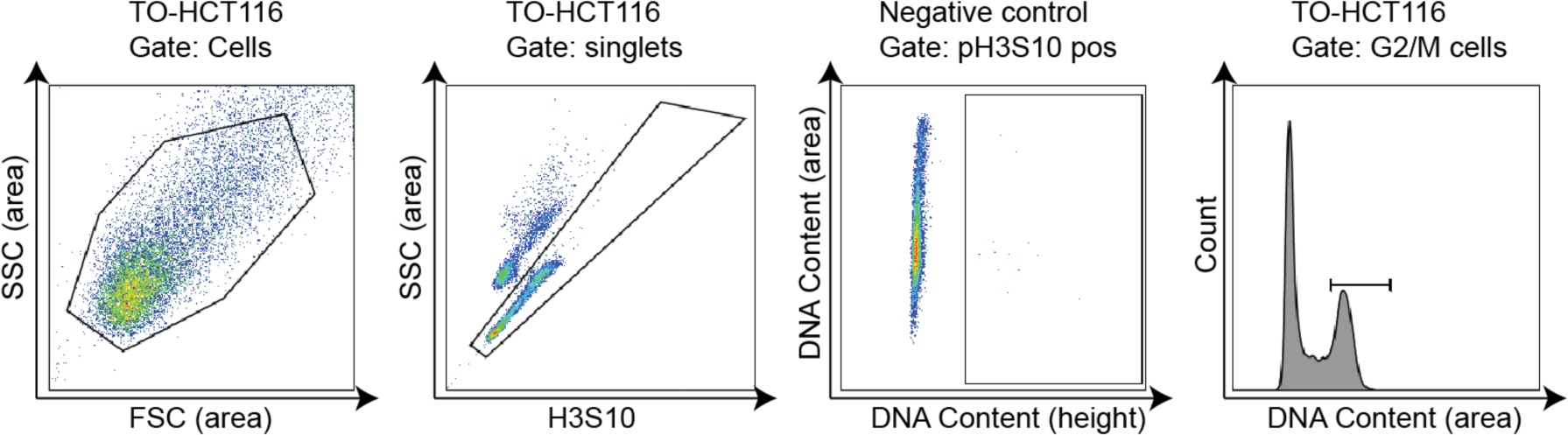
Flow cytometry gating. Example of flow cytometry gating for Figure 7a. FSC (area) vs SSC (area) gating was used to exclude the cell debris. Next, singlets were gated on FxCycle™ Violet (DNA content) height vs area. Phospho-H3S10 positive population was gated on DNA content (height) vs DNA content (area) of the unstained negative control cells. G2/M cell population was gated on the DNA content (area) histogram.

**Figure 8—Figure supplement 1.**
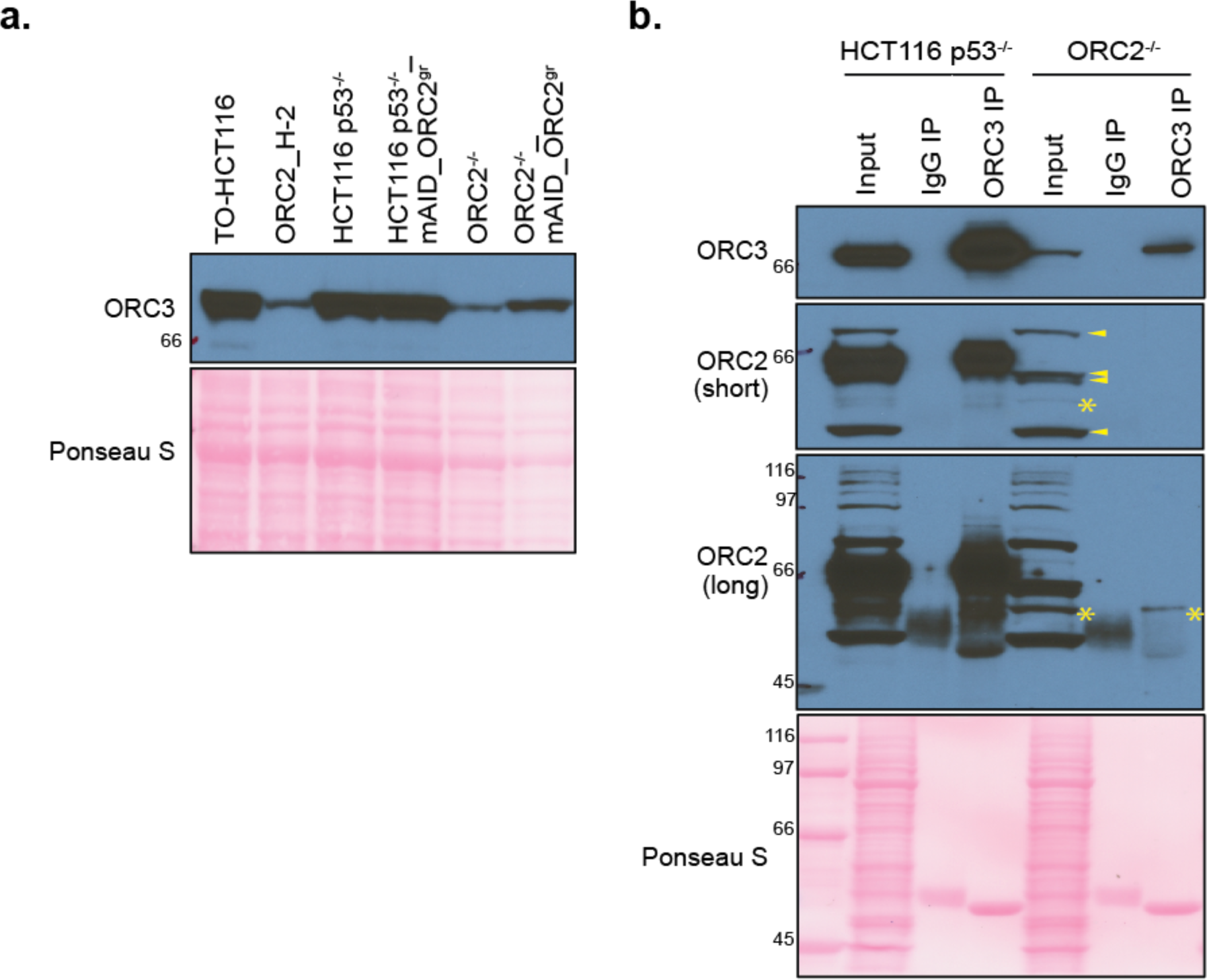
ORC3 exists in *ORC2^-/-^* cell line. (a) ORC3 expression in TO- HCT116, ORC2_H-2, HCT116 *p53^-/-^*, HCT116 *p53^-/-^*_mAID-ORC2^gr^, *ORC2^-/-^*, and *ORC2^-/-^* _mAID-ORC2^gr^ cell lines. Whole cells were boiled in Laemmli buffer and followed by western blotting and detected with anti-ORC3 antibody. (b) ORC3 immunoprecipitation (IP) in HCT116 *p53^-/-^* and *ORC2^-/-^* cell lines. Cells were lysed in lysis buffer and incubated with mouse IgG or ORC3 mouse monoclonal antibody for immunoprecipitation, followed by western blotting and detected with antibodies against ORC2 and ORC3. The loaded input was 2.5% and IP was 30%. Both short and long exposure of ORC2 detection were shown here. Asterisks (*) indicated the putative truncated ORC2 which was only found in *ORC2^-/-^* cell line. In the short exposure only, arrows pointed to nonspecific bands detected by the anti-ORC2 antibody.

**Figure 8—Figure supplement 2.**
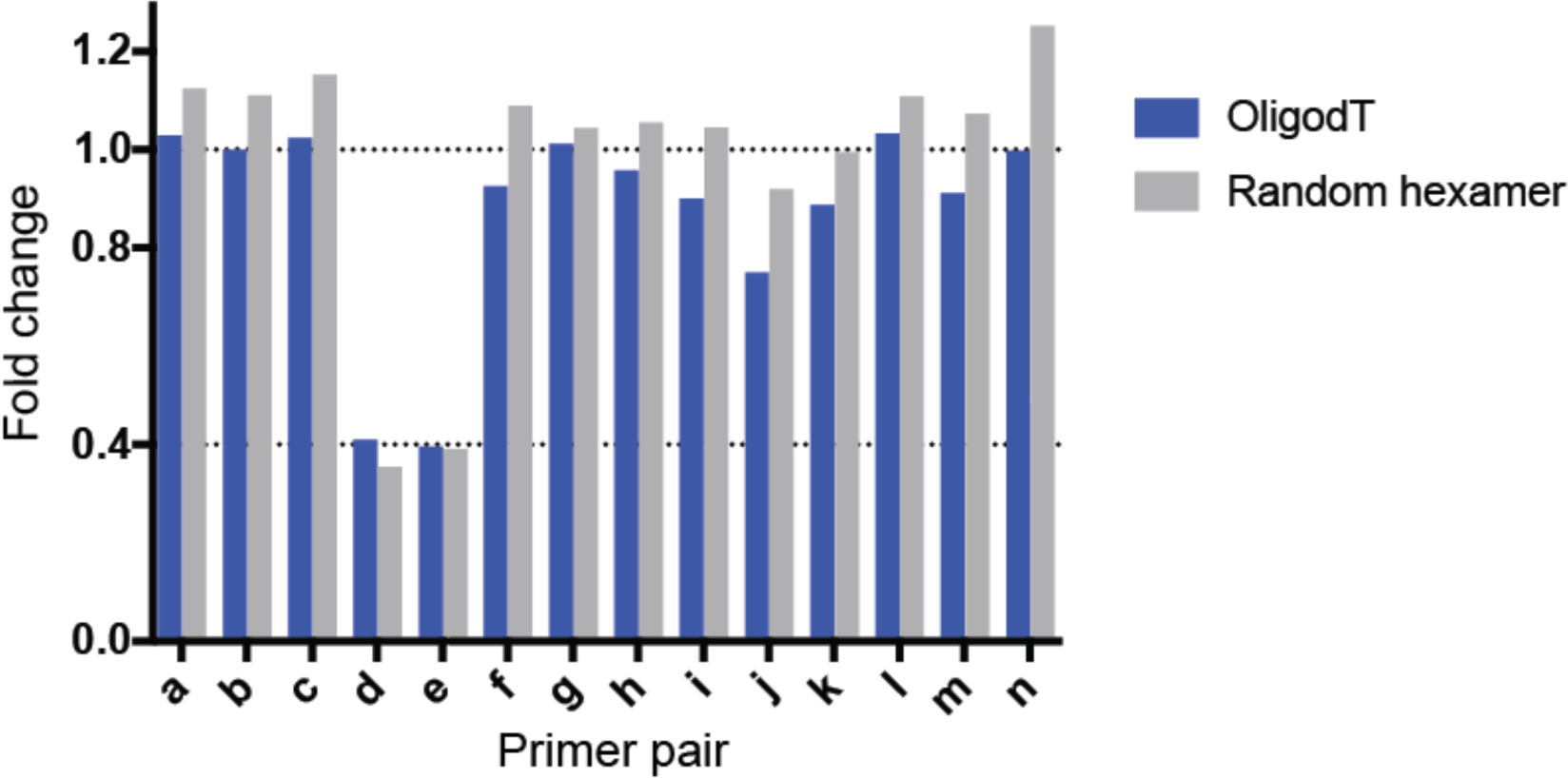
Real time quantitative PCR fold change in a bar diagram view. Blue, fold change of OligodT primer sample. Grey, fold change of random hexamer sample. The exon junctions in the *ORC2* cDNA are shown as indicated in Figure 8e and the Fold change (FC) for each primer pair in *ORC2^-/-^* cells compared to HCT116 *p53^-/-^* cells was calculated as FC = 2(to the power of ΔΔCt).

**Figure 8—Figure supplement 3.**
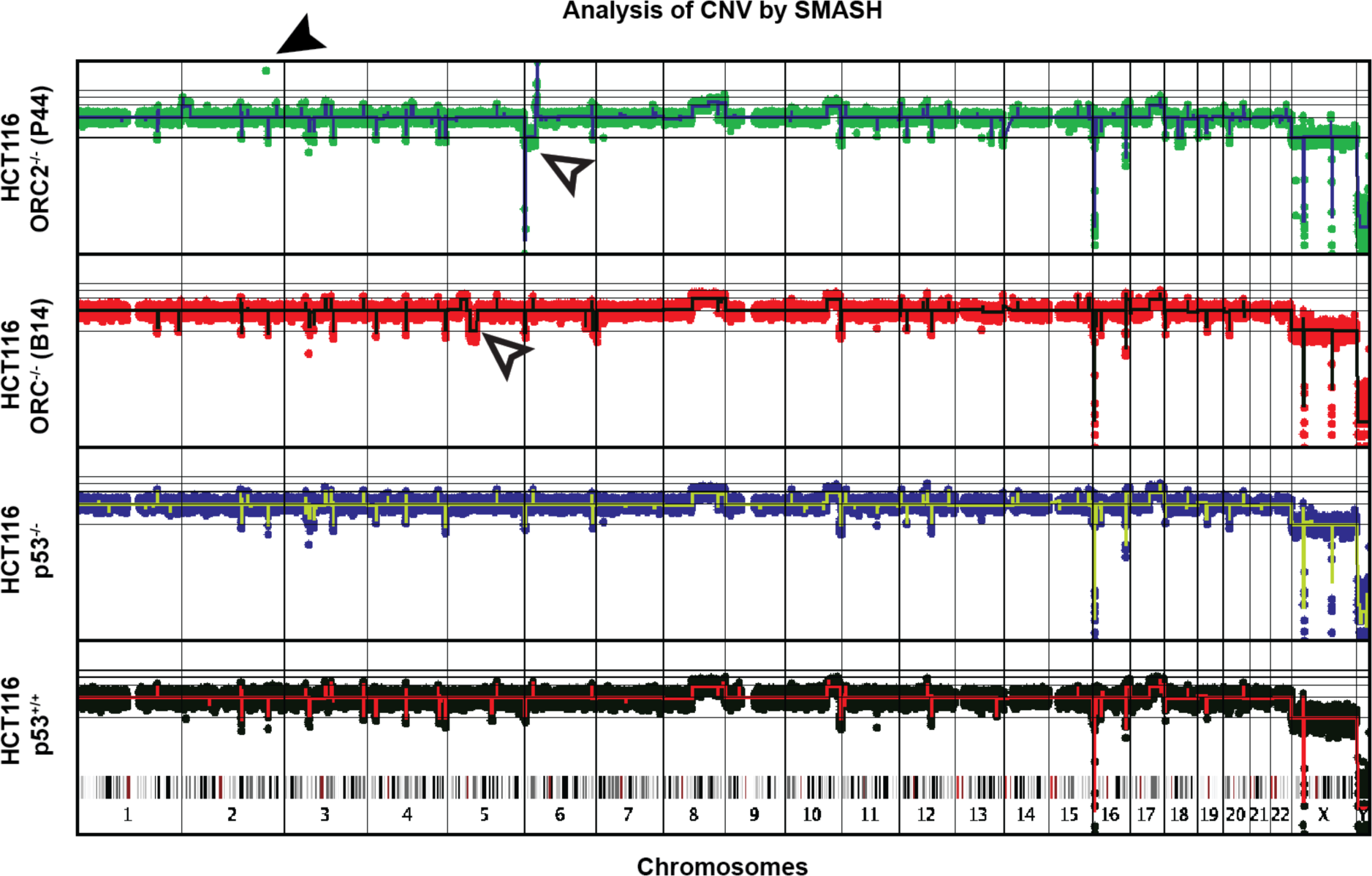
Copy number analysis of the genomes of four cell lines using the SMASH method. The amplification of the ORC2 gene sequences in HCT116 *ORC2^-/-^* (P44) cells is shown by the green dot and filled arrow. The open arrows show acquired CNVs in the *ORC1^-/-^* and *ORC2^-/-^* cells compared to the parent cells.

**Figure 8—Figure supplement 4.**
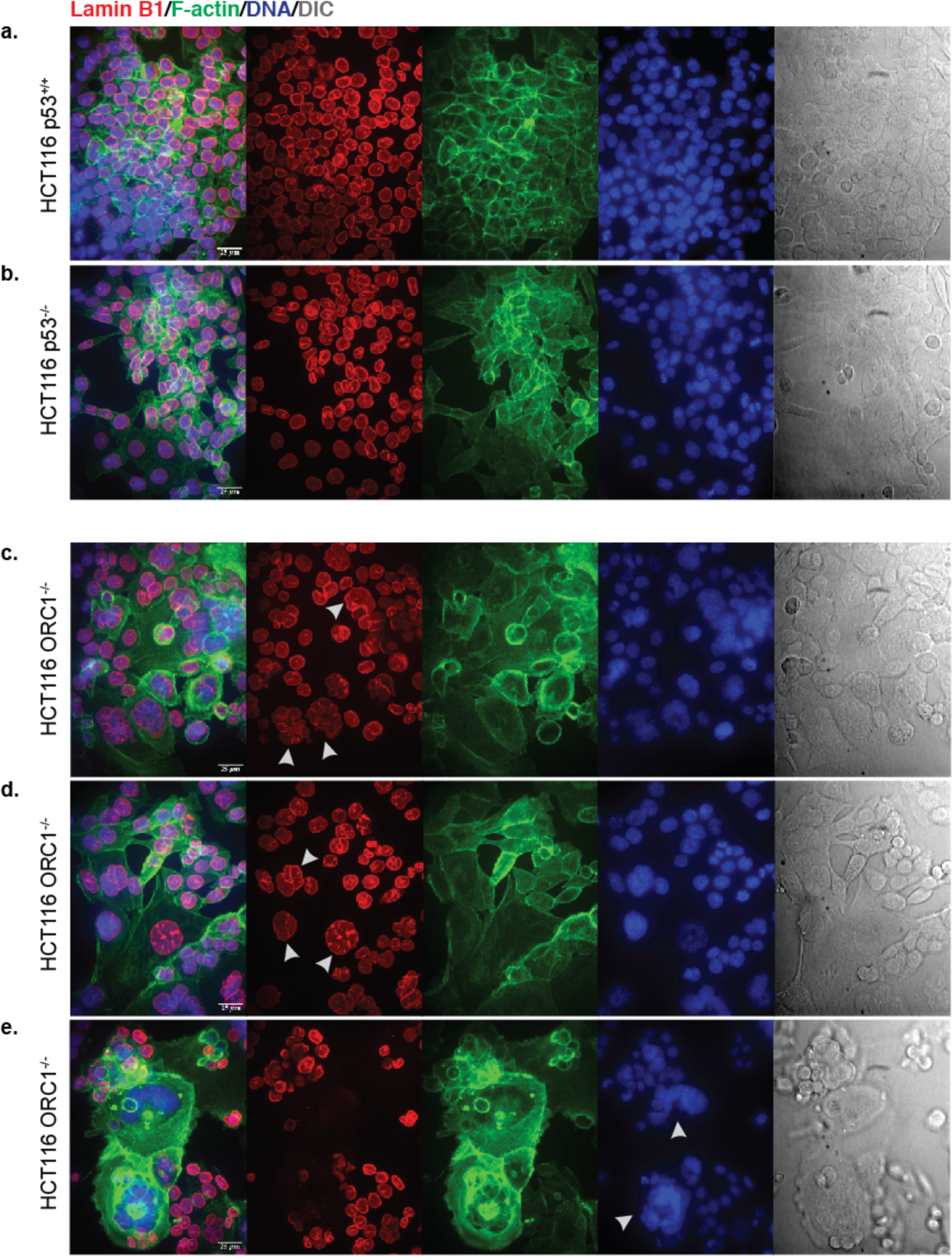
Confocal Microscopy images of HCT116 cell lines. Imaged acquired as z-stack of 25 µm (z = 1µm each). Images presented maximum intensity projections in the merge and average intensity projections in single channel images. Channel reference: Lamin B1 (Red), F-actin (Green), DNA (blue), DIC (grey) (a) HCT116 *p53^+/+^*. (b) HCT116 *p53 ^-/-^*. (c-e) HCT116 *ORC1^-/-^* (B14). Scale bar is 25 µm.

**8-Figure supplement 5.**
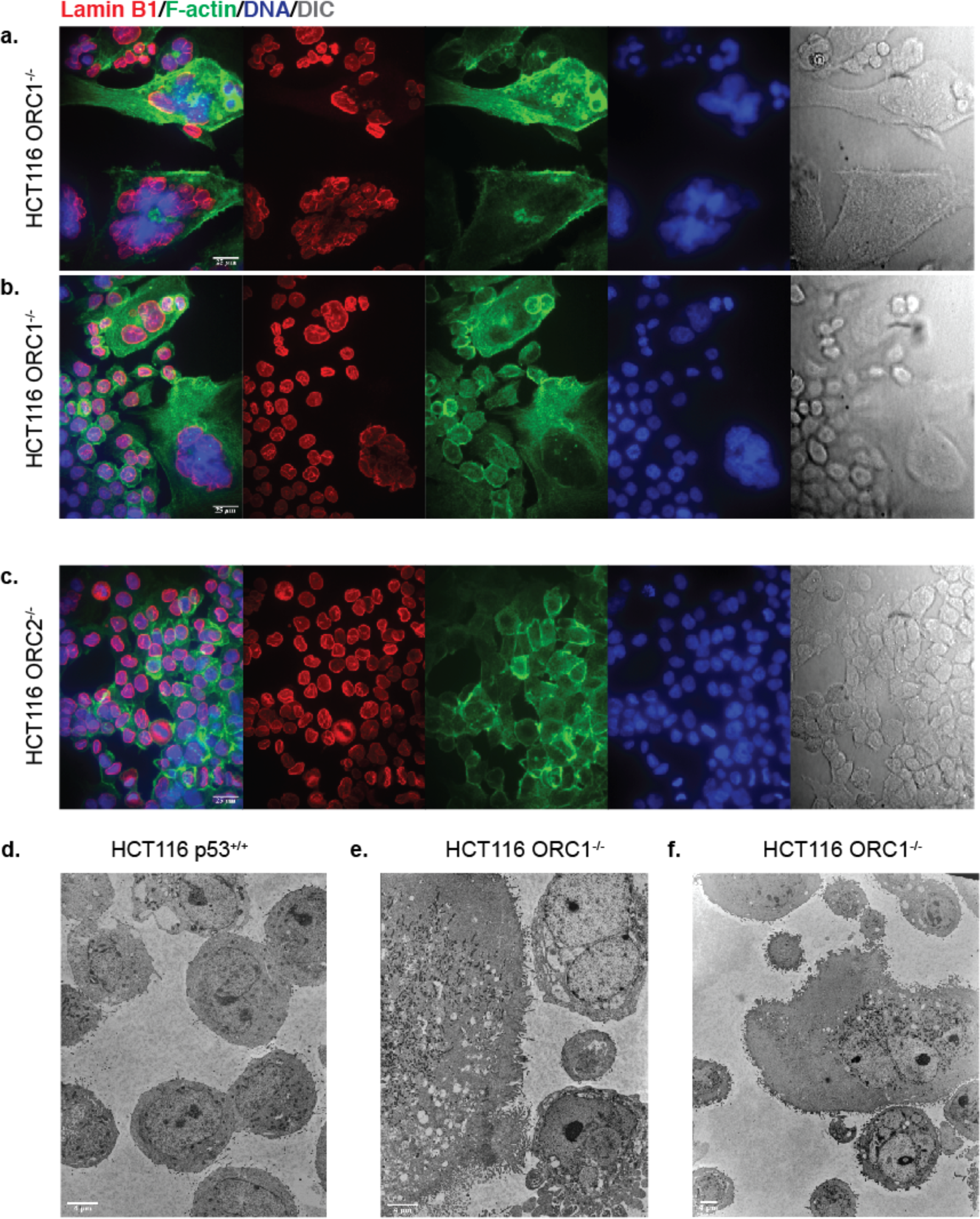
Confocal (a-c) and Transmission electron microscopy (d-f) images of HCT116 cell lines. Confocal: (a-b) HCT116 *ORC1·^1^* (814), (c) HCT116 ORC2· le bar is 25 IJm. TEM: (d) HCT116 *p53+^1^+* cells with 2000x magnification. (e) *ORC1·^1^* cells 000x magnification. (f) *ORC1·^1^* cells with 1000x magnification. Scale bar in (d-e) is 41 µm.

Supplement Table 1. The sequences of all guide RNAs used for gene editing, including those directed to ORC1-6 and CDC6 as well as positive and negative guides for the tiling CRISPR screens.

Supplement Table 2. Sequence of Barcode primers used for Next Gene Sequencing analysis in tiling CRISPR screens.

Supplement Table 3. Primers used for exon analysis qPCR of the *ORC2* gene cDNAs from various cell lines.

